# A historical cross-border Andes virus lineage reveals the origin of a cruise ship hantavirus pulmonary syndrome outbreak

**DOI:** 10.64898/2026.07.19.739363

**Authors:** Hade Ramos, Constanza Díaz-Gavidia, Dianne Díaz-Ramírez, Eugenia Fuentes-Luppichini, Jens H. Kuhn, Carla M. Bellomo, Andreas Schüller, Raúl Araya-Secchi, International Genomics Consortium Investigating the M/V Hondius Outbreak, Daniel M. Cisterna, Luciana Fernandez-Bettelli, Santiago Palacios-Aliggi, Valeria P. Martínez, Marcela Ferrés, Piet Maes, Gustavo Palacios, Nicole D. Tischler, Jenniffer Angulo

**Affiliations:** Departamento de Enfermedades Infecciosas e Inmunología Pediátricas, Escuela de Medicina, Pontificia Universidad Católica de Chile, Santiago, Chile; Facultad de Ciencias Biológicas, Pontificia Universidad Católica de Chile, Santiago, Chile; Escuela de Medicina Veterinaria, Facultad de Medicina, Pontificia Universidad Católica de Chile, Santiago, Chile; Facultad de Agronomía y Sistemas Naturales, Pontificia Universidad Católica de Chile, Santiago, Chile; Laboratorio de Virología Molecular, Centro Ciencia & Vida, Fundación Ciencia & Vida,, Santiago, Chile; Doctorado en Biotecnología y Bioemprendimiento, Facultad de Medicina, Universidad San Sebastián, Santiago, Chile; Department of Microbiology, Icahn School of Medicine at Mount Sinai, New York, NY, USA; Laboratorio Nacional de Referencia de Hantavirus, Instituto Nacional de Enfermedades Infecciosas, Administración Nacional de Laboratorios e Institutos de Salud Dr. Carlos G. Malbrán; Buenos Aires, Argentina; Institute for Biological and Medical Engineering, Pontificia Universidad Católica de Chile, Santiago, Chile; Facultad de Ingeniería, Universidad San Sebastián, Valdivia, Chile and Centro de Estudios Científicos, Valdivia, Chile; The full list of the authors and affiliations of this consortium can be found in the Supplementary information; Instituto Nacional de Enfermedades Infecciosas, Administración Nacional de Laboratorios e Institutos de Salud Dr. Carlos G. Malbrán; Buenos Aires, Argentina; Laboratorio de Infectología y Virología Molecular, Red Salud UC-Christus, Santiago, Chile; SENTINET, Santiago, Chile; European Plotkin Institute for Vaccinology, Université Libre de Bruxelles, Brussels, Belgium; Global Health Emerging Pathogens Institute, Icahn School of Medicine at Mount Sinai, New York, NY, USA; Escuela de Bioquímica, Facultad de Ciencias, Universidad San Sebastián, Santiago, Chile

**Author notes:** These authors contributed equally to this work (Hade Ramos, Constanza Díaz-Gavidia, Dianne Díaz-Ramírez, Eugenia Fuentes-Luppichini, Jens H. Kuhn).

## Abstract

Andes virus (ANDV) caused a multi-country outbreak of hantavirus pulmonary syndrome among passengers and crew of a cruise ship in 2026. To investigate the origin and evolutionary history of the virus responsible for the outbreak, we analyzed complete ANDV small (S), medium (M), and large (L) genome segment sequences from Chile alongside outbreak-associated and publicly available genome sequences. Across all three segment-specific phylogenies, the outbreak virus clustered within an ANDV Clade III cluster spanning southern Chile and northern Patagonia in Argentina and were most closely related to a human-derived ANDV (p1236) collected in Los Ríos Region of Chile in 2012, representing the closest known historical relative of the outbreak-associated virus. Phylogeographic analysis showed that the genetically distinct ANDV Clade V lineage circulating in central Chile was not closely related to the cruise ship outbreak-associated genomes, thereby reducing the likelihood that the index cases acquired infection through zoonotic spillover while traveling through the Maule Region. These findings trace the geographic origin of the outbreak-associated virus to a defined corridor, the Hua Hum Pass, a cross-border zone connecting Province of Neuquén in Argentina with the Los Ríos and La Araucanía Regions of Chile.

## Introduction

The April–May 2026 outbreak of hantavirus pulmonary syndrome (HPS) aboard the Dutch-flagged *M/V Hondius* cruise ship^1^ represented an unusual international displacement of an orthohantavirus, apparently initiated by zoonotic spillover and followed by human-to-human transmission during the ship voyage. Recent genomic and epidemiological investigations characterized the outbreak variant and reconstructed key elements of the travel history of the presumed index cases^1^. However, the precise geographic site and circumstances of the initial zoonotic spillover remain unresolved.

Andes virus (ANDV) is a rodent-borne orthohantavirus in subfamily *Mammantavirinae* of family *Hantaviridae*. It is classified within the species *Orthohantavirus andesense* together with three closely related “ANDV-like” viruses, Castelo dos Sonhos virus (CASV), Lechiguanas virus (LECV), and Orán virus (ORNV). Like all orthohantaviruses, ANDV has a trisegmented negative-sense RNA genome^2^.

Genomic analysis identified the cruise ship outbreak agent as ANDV and showed that the outbreak-associated genome sequences were most closely related to ANDV genome sequences causing HPS cases in Province of Neuquén, Argentina^3,4^. In particular, the cruise ship outbreak virus genome sequence clustered with sequences from HPS cases in Villa Meliquina, San Martín de los Andes, and Villa Traful, documented from 2017 to 2026^4^.

Pairwise comparisons showed approximately 98.7% nucleotide identity between the cruise-associated virus sequence and the closest Province of Neuquén small (S), medium (M), and large (L) genome segment sequences^4^. The cruise variant reference sequence, ANDV-Switzerland-Hu-3337-2026, differs from the NRC4/2018 Province of Neuquén reference sequence by 23 single nucleotide variants (SNVs) in the S, 47 in the M, and 84 in the L segment. Most mutations were synonymous; nonsynonymous mutations were predicted to result in V70I and K144R changes in the large (L) protein (encoded by the L segment), and T193A, T516I, and A994T changes in the glycoprotein precursor (GPC; encoded by the M segment)^3,5^. These genome sequences from Province of Neuquén were interpreted^3,4^ as the closest sampled human representatives of a broader ANDV population circulating across northern Patagonia in Argentina and southern Chile. However, they are not sufficiently closely related to be considered direct ancestors of the cruise ship outbreak-associated virus.

Closely related rodent-derived sequences were recently made available^6^. In a pairwise comparison against the Swiss cruise-ship reference genome sequence ANDV/Switzerland/Hu-3337/2026 (PZ385161.1, PZ385162.1, and PZ385163.1)^7^, the closest rodent-derived ANDV sequence from Chile in the available set, R175, still differs by 68 unambiguous nucleotide substitutions across the three segments: 36 SNVs in L, 23 in M, and 9 in S. This level of divergence is still substantially greater than that reported by Kim et al.^8^, who used active targeted surveillance to identify the geographical site of emergence of a distantly related orthohantavirus, Hantaan virus. In their study, paired human and rodent Hantaan virus sequences differed by only 8–30 nucleotide substitutions across the complete trisegmented genome, depending on the human–rodent pair. Thus, the currently available rodent sequences from Chile should be interpreted as evidence of broader unsampled regional diversity rather than as the likely immediate source.

Our epidemiological reconstruction showed that the presumed index cases, Cases 1 and 2, travelled extensively by motorhome through Argentina, Uruguay, and Chile before embarking on the cruise in Ushuaia, Province of Tierra del Fuego, Antarctica and South Atlantic Islands, Argentina, on 1 April 2026. This work will focus on their travel through Chile. They entered Chile’s Aysén Region on 9 January, returning to Argentina via the Icalma Pass on 31 January. These regions are known for the circulation of ANDV and the presence of long-tailed pygmy rice rats (*Oligoryzomys longicaudatus* (E. T. Bennett, 1832)), ANDV’s main spillover rodent reservoirs^9^. On 12 February, Cases 1 and 2 moved via the Pehuenche Pass to Maule Region in Chile, and re-entered Argentina four days later via the same pass.

Recently generated complete genome sequence data from human-derived ANDV in Chile, spanning a period between 2011–2024^10^, now provides an opportunity to evaluate this possibility in a higher-resolution phylogeographic framework. That dataset defines geographically structured ANDV diversity in Chile, including a new northern/central lineage (ANDV Chi-North, from now on referred to as ANDV Clade V). In the present study, we expanded this framework by incorporating additional 2025–2026 sequences from Chile together with HPS outbreak-associated ANDV genomes. By comparing the cruise ship outbreak variant sequence with this expanded dataset and by mapping those relationships against the reconstructed travel itinerary of the index cases, we evaluate whether the initial zoonotic ANDV spillover could have occurred in Chile or in an unsampled cross-border extension of the northern Patagonia from Argentina.

## Results

### Cruise ship outbreak-associated Andes virus genome sequences cluster with those of Andes viruses circulating in the Hua Hum Pass corridor

To investigate the ancestry of the cruise ship-associated ANDV, we leveraged three sequence datasets populated exclusively with complete genome sequences comprising the ANDV S, M, and L segments: cruise ship outbreak-associated ANDV, Argentina-endemic ANDV, and Chile-endemic ANDV genome sequences. This approach minimized biases arising from segment-specific phylogenetic discordance while enabling direct comparison across all three genome segments. Maximum-likelihood phylogenetic analyses placed each cruise ship outbreak-associated genome segment sequence within previously described ANDV diversity, clustering with sequences belonging to ANDV Clade III^2^, in a distinct cluster, here designated “Hua Hum Pass corridor (HHPC)”, composed predominantly of historical human-derived ANDV genome sequences from La Araucanía and Los Ríos Regions in Chile together with related northern Patagonian sequences from Argentina (**Fig. 1**).

**Fig. 1.**
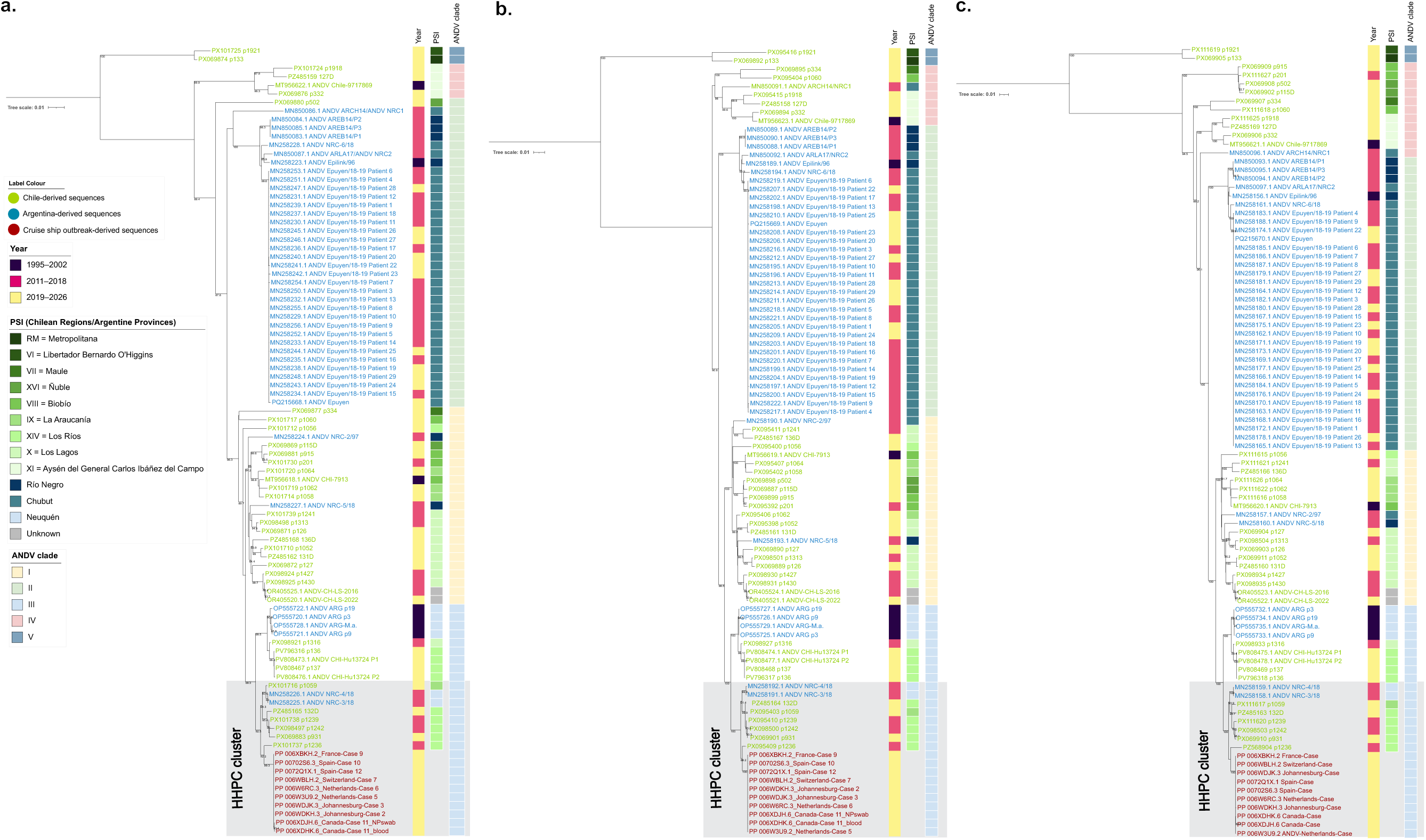
Cruise ship outbreak-associated Andes virus genome sequences cluster with those of Andes viruses circulating in the Hua Hum Pass corridor. Maximum-likelihood (ML) phylogenetic trees were reconstructed independently for the complete (**a**) S, (**b**) M, and (**c**) L genomic segment sequences using cruise ship outbreak-associated Andes virus (ANDV) genome sequences together with representative ANDV reference sequences from Chile and Argentina. Clade denomination follows Bellomo et al.^11^. Branch support values correspond to SH-aLRT support obtained from 1,000 replicates. Trees were reconstructed using IQ-TREE and visualized with iTOL.

Within the S segment phylogeny (**Fig. 1a**), cruise ship outbreak-associated genome sequences clustered with sequences derived from Chile p1236 (Los Ríos Region, 2012), p931 (Los Ríos Region, 2020), p1242 (Los Ríos Region, 2013), p1239 (Los Ríos Region, 2013), p132D (Los Ríos Region, 2026), and p1059 (La Araucanía Region, 2023), together with Argentinian Province of Neuquén genome sequences NRC-3/18 and NRC-4/18. Similar clustering patterns were observed independently in the M (**Fig. 1b**) and L (**Fig. 1c**) segment phylogeny.

Bayesian phylogenetic analyses recovered highly consistent maximum clade credibility (MCC) trees, corroborating the phylogenetic placement inferred by maximum-likelihood analyses (**Supplementary Figure S1**).

Within the HHPC cluster, p1236 emerged as the closest sampled human-derived relative of the cruise ship outbreak-associated viruses across all three genome segments. The patient was a 7-year-old male from the Lanco commune (Los Ríos Region, southern Chile) who developed HPS following an isolated rural exposure, with no associated cluster or secondary cases. He was treated at Panguipulli Hospital, located within the Hua Hum Pass corridor identified in our phylogeographic analyses, where he developed severe respiratory failure requiring invasive mechanical ventilation and subsequently recovered. Although the p1236 L segment contained two unresolved gaps (positions 4008–5185 and 5461–6562), the resolved sequence supported the same phylogenetic placement observed for the S and M segments, with p1236 consistently clustering more closely with the cruise ship outbreak-associated genomes than NRC-4/2018^3,4^. Direct comparison between p1236 and the Swiss outbreak-associated genome sequence identified 18 nucleotide differences in the S segment (no nonsynonymous substitutions), 28 in the M segment (two nonsynonymous substitutions), and 78 across the 4,535-nt available portion of the L segment (three nonsynonymous substitutions). Together, these findings identify a historical human-derived ANDV genome from southern Chile that is more closely related to the outbreak-associated viruses than the previously available genome sequences from Argentina.

Motivated by published evidence suggesting potential reassortment including the phylogenetic placement of the publicly available rodent-derived genome sequence R600 from the Los Ríos Region of Chile (2013), which clustered with the outbreak-associated genome sequences in the L-segment phylogeny but not in the S and M phylogenies^7^, we performed a comprehensive reassortment analysis across our complete dataset. This analysis detected little evidence of reassortment among Clade III genomes (**Supplementary Figure S2**), supporting the interpretation that the HHPC cluster represents a single evolutionary lineage.

Several additional human-associated genome sequences composing the HHPC cluster were obtained from HPS cases identified since 2012 in southern Chile. These cases included sporadic infections, epidemiologically linked clusters, and a range of clinical outcomes (**Supplementary Table S1)**. These findings indicate that the HHPC cluster represents a long-standing component of ANDV diversity in southern Chile and northern Patagonia in Argentina.

### The Hua Hum Pass corridor Andes virus cluster is defined by limited, largely neutral molecular signatures within Andes virus Clade III diversity

No major genomic insertions, deletions, or segment-specific genomic features distinguished HHPC cluster genome sequences from those of other ANDV Clade III members.

Comparison of deduced amino acid sequences identified few phylogenetically informative substitutions. Consistent with previous analyses^10^, ANDV Clade III was characterized by substitutions including GPC I114V, A193T and V1127I. Within this background, genomes composing the HHPC cluster consistently encoded GPC T1055, whereas the remaining analyzed ANDV Clade III genomes encoded GPC A1055. Additional cluster-enriched changes included nucleoprotein (S segment) substitution L7I and L protein (L segment) R144K and A541V. These substitutions define a molecular signature that distinguishes the HHPC cluster within the broader diversity of ANDV Clade III (**Supplementary Table S2**). A comprehensive comparison of all deduced amino acid substitutions identified among HHPC cluster genome sequences relative to the five recognized ANDV clades (Clades I–V) and representative orthohantaviruses is provided in **Supplementary Table S3**.

Previously proposed amino acid markers for person-to-person transmission in Argentinian HPS outbreaks^11^ were not consistently encoded in HHPC cluster genome sequences. The only substitution unique to the cruise ship outbreak lineage was GPC T516I, which remained exclusive after inclusion of the expanded genomic dataset from Chile.

Structural mapping of HHPC cluster-associated aminoacidic substitutions onto the GPC-encoded G_N_/G_c_ complex showed that three of the five substitutions are located in surface-exposed regions of the spikes, whereas the remaining two mapped to the G_N_ endodomain and G_C_ transmembrane region (**Supplementary Figure S3**). None of the surface-exposed substitutions overlapped with previously reported neutralizing antibody epitopes or escape variants. Similarly, the three L protein substitutions mapped outside of the known catalytic sites of this multifunctional protein^12^ (**Supplementary Figure S4**). Deep learning-based structural predictions indicated only minor changes in Gibbs free energy (ΔΔG), suggesting minimal effects on protein stability or quaternary assembly. Together, these findings suggest that the HHPC-associated substitutions represent geographically localized signatures rather than adaptive changes affecting viral fitness.

### Phylogeographic network analysis identifies circulation of Hua Hum Pass corridor Andes viruses in an area that links La Araucanía/Los Ríos Regions in Chile with Province of Neuquén in Argentina

Geographic mapping showed a broad distribution of Clade III ANDV across endemic areas of southern Chile and Argentinean northern Patagonia (**Fig. 2a**). Colored circles represent geographically clustered ANDV genome sequences included in the present study together with previously reported human-derived genomes from Chile^10^. Cruise ship outbreak-associated genome sequences clustered within a HHPC cluster distributed primarily across La Araucanía, Los Ríos, and Los Lagos Regions in southern Chile, whereas related genome sequences from Argentina were distributed across northern Patagonian localities including Epuyén and El Hoyo in Province of Chubut, Villa Meliquina in Province of Neuquén, and El Bolsón and Bariloche in Province of Río Negro.

**Fig. 2.**
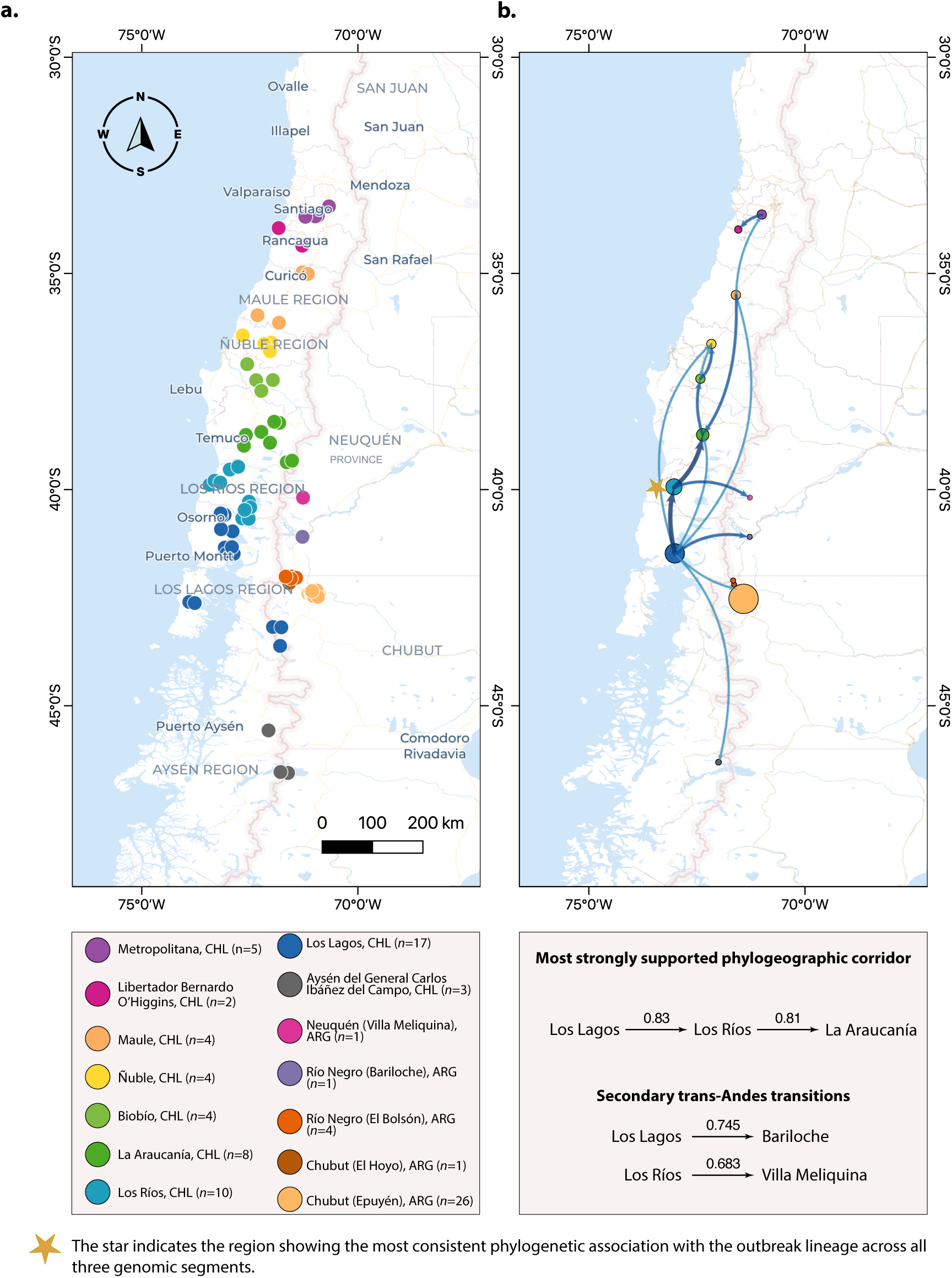
Phylogeographic network analysis identifies circulation of Hua Hum Pass corridor Andes viruses in an area that links La Araucanía/Los Ríos Regions in Chile with Province of Neuquén in Argentina. (**a**) Geographic association of Andes virus (ANDV) genome sequences included in the phylogenetic analyses. Each colored circle represents the probable site of infection of hantavirus pulmonary syndrome case from which an ANDV genome sequence was determined. The dataset comprises all human-derived ANDV genome sequences from Chile with available epidemiological information reported in^10^ together with publicly available genome sequences from Argentina (Fig. 1) and are grouped according to geographic clustering patterns. (**b**) Phylogeographic transition corridors inferred from Bayesian discrete trait analyses among ANDV-endemic regions. Arrows indicate the most strongly supported transitions between Chilean and Argentinian localities. Arrow width and color intensity are proportional to posterior transition support values.

Phylogeographic reconstruction inferred supported transition corridors between endemic regions in southern Chile and northern Patagonia in Argentina (**Fig. 2b**). Directional arrows indicate inferred transitions between geographic regions. The strongest supported transitions connected Los Lagos, Los Ríos, and La Araucanía Regions in southern Chile, whereas additional trans-Andean transitions connected Chilean Los Lagos Region with Argentinian Province of Río Negro and Chilean Los Ríos Region with Argentinian Province of Neuquén. This area is anchored by the Hua Hum Pass, which connects Panguipulli/Puerto Fuy in Los Ríos Region, Chile, with San Martín de los Andes, Province of Neuquén, in Argentina. This section of the Andes is characterized by one of the lowest elevations along the Chilean Andes (approximately 685 m above sea level), facilitating ecological connectivity between southern Chile and Argentinian’s Province of Neuquén.

For instance, the Hua Hum Pass provides a geographic link between the phylogeographic reconstruction and the travel history of the presumed index cases.

### Cruise ship outbreak-associated index cases land travel route included regions where Andes viruses phylogenetically related to cruise ship Andes viruses are endemic

Integration of the reconstructed land travel itinerary before cruise ship embarkation with the phylogenetic and phylogeographic analyses revealed extensive geographic overlap between the movements of index Cases 1 and 2 and regions where HHPC cluster ANDV has been identified (**Fig. 3**). From 7 to 31 January 2026, Cases 1 and 2 travelled extensively through southern Chile. They entered Chile from Argentina via Jeinimeni Pass on 7 January and remained in Aysén Region through 13 January, travelling through Chile Chico, Río Ibáñez, Coyhaique, Lago Verde, and Cisnes. On 14 January, they entered Los Lagos Region, where they travelled through Chaitén, Hualaihué, Ancud (Chiloé), and Dalcahue until 26 January. They entered Los Ríos Region on 27 January, visiting Los Lagos and Mariquina, followed by Araucanía Region on 28 January, where they travelled through Toltén and Lonquimay until 31 January. Their itinerary concluded on 31 January with travel from Araucanía into Province of Neuquén, Argentina, near Aluminé. However, our recent nationwide genomic characterization showed that ANDV diversity in Chile is strictly compartmentalized by geography^10^. Clade III, including the HHPC cluster, maps to the areas visited by the index cases between 27 and 31 January, whereas Clade II, including the Epuyén/18 and Epilink/96 variants associated with enhanced person-to-person transmission ability^13^, was not detected in Chile. Notably, the index cases visited Los Lagos and Mariquina in Los Ríos Region, within the area where p1236 was detected in Lanco in 2012. Although the itinerary does not include Lanco specifically, this regional overlap links their travel route to the known distribution of the HHPC cluster.

**Fig. 3.**
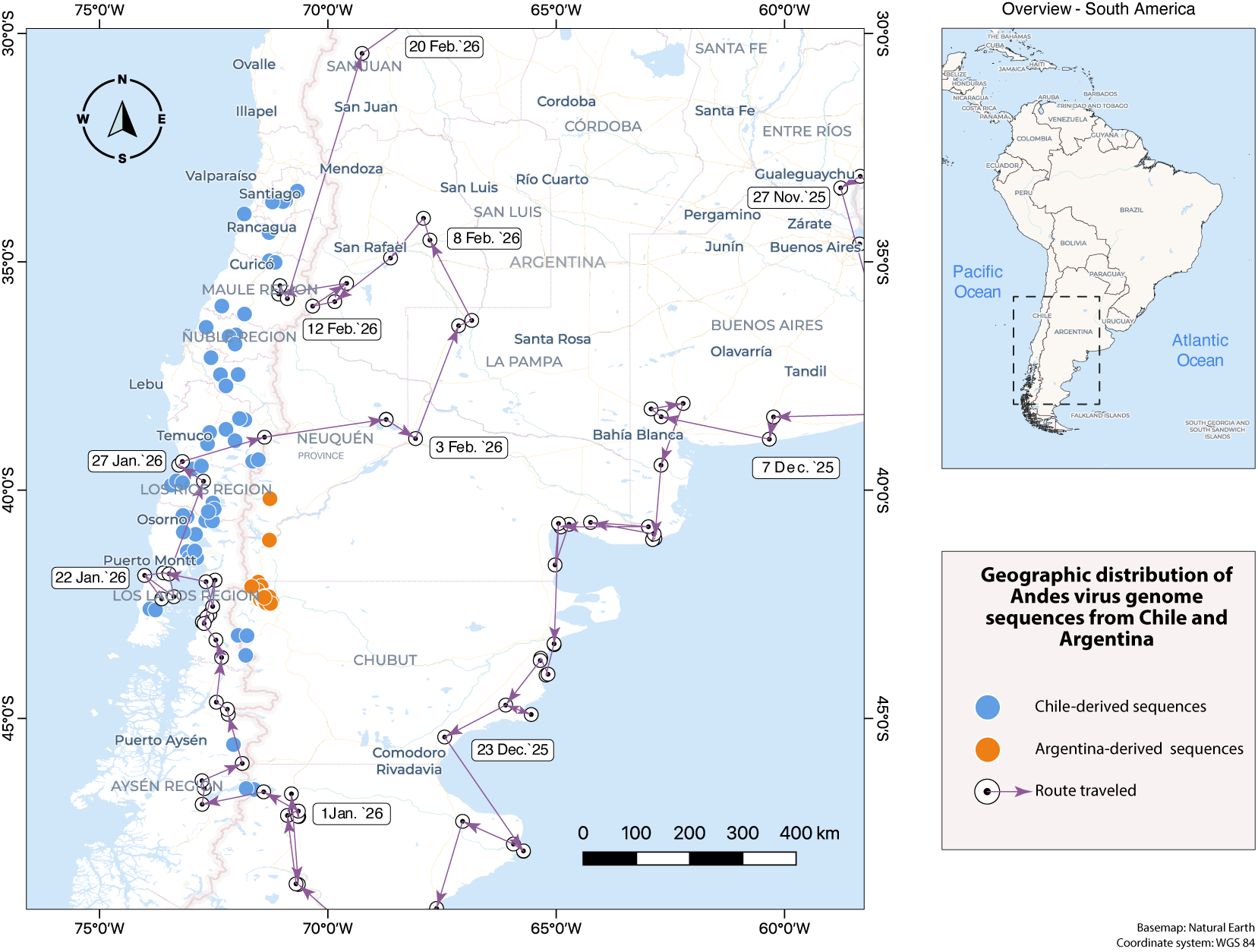
Cruise ship outbreak-associated index case land travel route included regions where Andes viruses phylogenetically related to cruise ship Andes viruses are endemic. Geographic reconstruction of the pre-embarkation land travel itinerary of the presumed cruise ship outbreak index Case 1 and 2 before boarding the cruise ship. The travel route is shown as directional arrows connecting documented travel localities reconstructed from epidemiological investigation records. Colored circles indicate the probable sites of infection of later hantavirus pulmonary syndrome cases from whom Andes virus (ANDV) genome sequences were determined, comprising ANDV genome sequences from Chile^10^ and Argentina. Inset: overview map of South America showing the location of the study area.

On the other hand, the itinerary also included travel through Maule Region from 12 to 16 February 2026, an area corresponding to the geographic range of the genetically distinct, Clade V ANDV^10^. However, the outbreak-associated genome sequences have no phylogenetic relationship with ANDV Clade V genome sequences.

Taken together, these observations indicate that the travel route intersected regions associated with ANDV of Clades III and V in Chile, whereas the outbreak-associated genome sequences remained phylogenetically linked to Clade III viruses circulating across the cross-border of southern Chile and northern Patagonia from Argentina.

## Discussion

The cruise ship HPS outbreak represents an unusual international displacement of ANDV, likely initiated by zoonotic infection before embarkation and followed by person-to-person transmission in a confined travel setting. Although previous epidemiological and genomic investigations established that the outbreak virus belonged to ANDV Clade III and supported onward transmission aboard the vessel^3^, the location of the initial spillover event remained unresolved.

By incorporating newly generated complete genome sequences of ANDV from Chile, we substantially expanded the phylogenetic context of the outbreak and provide strong evidence that the outbreak virus emerged from a long-standing transboundary population circulating across southern Chile and norther Patagonia in Argentina for more than a decade, rather than from a recently emerged variant.

Across all three genome segments, cruise ship outbreak-associated genomes consistently clustered within the HHPC cluster of ANDV Clade III, which includes historical human-derived genomes from La Araucanía and Los Ríos Regions in Chile together with related genomes from Neuquén Province, Argentina. Available human- and rodent-derived ANDV genomes define the broader regional ancestry of the outbreak virus but do not identify the immediate reservoir population from which spillover occurred.

The reconstructed itinerary of the presumed index cases is consistent with this phylogeographic framework. Between 7 and 31 January 2026, before entering Argentina, they travelled through Los Lagos, Los Ríos, and La Araucanía Regions, overlapping the known distribution of the HHPC cluster and the inferred trans-Andean transition corridors linking southern Chile with Argentinian Provinces of Neuquén, Río Negro, and Chubut.

The itinerary also included travel through Chile’s Maule Region and Argentina’s Province of Mendoza from 12 to 16 February 2026, a period compatible with the reported HPS incubation period (approximately 43–45 days)^14^. However, genomic evidence argues against this scenario, as the outbreak-associated genomes showed no phylogenetic affinity with the genetically distinct ANDV Clade V circulating in central Chile. Furthermore, current ecological and epidemiological data do not document ANDV circulation in Province of Mendoza. Nevertheless, spillover during this leg of the journey cannot be excluded because of the limited genomic data available from local rodent populations.

Yet this interpretation introduces a new epidemiological challenge: the presence of the presumed index cases in Los Ríos Region on 27 January 2026, where the closest human-derived ANDV relatives of the cruise ship outbreak virus were identified, would imply an incubation period of approximately 66 days before symptom onset on 3 April 2026, substantially exceeding the currently accepted upper range for ANDV infection^14^. Thus, rather than resolving the origin of the outbreak, the expanded genomic dataset narrows its most likely geographic source while uncertainty of the precise timing and circumstances of spillover remain.

One plausible explanation reconciling the phylogeographic inference with the epidemiological timeline is delayed re-exposure to virus-contaminated materials following the initial travel period, such as contamination of the motorhome while traveling through the Hua Hum Pass corridor and subsequent aerosolization during cleaning or handling shortly before cruise embarkation. This hypothesis is compatible with the known environmental stability of orthohantaviruses^15^ and established incubation periods, although it remains speculative and requires epidemiological confirmation. Alternative explanations remain possible, including prolonged environmental persistence, incomplete reconstruction of the exposure history, an unidentified intermediate human infection, an unusually extended incubation period, or an unknown rodent spillover source host. None of these possibilities can currently be distinguished based on the available epidemiological and genomic data, highlighting the need for further epidemiological investigation.

Several uncertainties remain. ANDV sequence availability remains uneven across Chile and Argentina, particularly for rodent-derived viruses and regions traversed by the presumed index cases. Exact exposure sites and activities could not be fully reconstructed, limiting the ability to directly link phylogenetic findings with specific environmental exposures. In addition, phylogeographic transition estimates describe connectivity among sampled ANDV populations rather than direct evidence of recent rodent movement or human exposure. Finally, the cruise ship outbreak virus may have originated from an unsampled rodent reservoir population, particularly in trans-Andean regions where viral circulation likely extends beyond administrative boundaries. Rather than identifying a single source locality, our findings define the principal surveillance gap: coordinated, geographically dense genomic surveillance of both human HPS cases and rodent reservoir populations across the trans-Andean distribution of the HHPC cluster.

Our findings also provide new insight into the evolutionary history of the HHPC cluster. Identification of p1236 as the closest sampled human-derived relative of the cruise ship outbreak-associated virus across all three genome segments demonstrates that closely related ANDV diversity was already circulating in southern Chile in 2012. This finding further suggests that additional ANDV diversity remains unsampled across southern Chile and adjacent northern Patagonia, highlighting the importance of complete trisegmented genome sequencing for reconstructing ANDV evolutionary history and phylogeographic relationships.

Although pediatric HPS cases are less frequently reported than adult cases, they constitute a recognized component of ANDV epidemiology in Chile and may include severe clinical presentations^16^. The pediatric origin of p1236 therefore adds a valuable historical genome from an underrepresented patient group.

The HHPC cluster shows limited molecular divergence from the broader diversity of ANDV Clade III, with GPC S1055T representing its principal defining signature. Comparative genomic and structural analyses provided no evidence of major genomic or protein-level changes supporting the emergence of a uniquely adapted variant. Likewise, amino acid markers previously proposed to be associated with person-to-person transmission in Argentinian HPS outbreaks ^13^ were not consistently encoded by HHPC genomes. These observations indicate that the molecular characteristics of the HHPC cluster primarily reflect long-term regional diversification rather than recent adaptive evolution. Although GPC T516I was unique to the cruise ship outbreak-associated genomes, structural analyses predicted no measurable effects on glycoprotein stability, supporting its interpretation as a molecular marker of the common-source outbreak rather than a phenotypic determinant. Experimental studies are needed to determine whether T516I or other HHPC-associated substitutions have functional consequences.

Future surveillance should prioritize La Araucanía, Los Ríos, and Los Lagos Regions in Chile, and Provinces of Neuquén, Río Negro, and Chubut in Argentina, particularly along travel corridors, border crossings, and areas where human exposure to long-tailed pygmy rice rats may occur. The identification of related genome sequences spanning more than a decade also emphasizes the value of integrating historical genomic data into outbreak investigations. Continued full-genome sequence surveillance combining human, rodent, ecological, and geographic data will be essential to resolve future spillover events and to better characterize the ecological and evolutionary processes shaping ANDV diversity and transmission.

## Online material and methods

### Study overview and datasets

This study was conducted as part of an international collaborative investigation integrating genomic and epidemiological data associated with the 2026 cruise ship hantavirus pulmonary syndrome (HPS) outbreak in southern South America. Comparative phylogenetic and genomic analyses were performed to investigate the evolutionary placement and regional context of outbreak-associated Andes virus (ANDV) genome sequences.

Three datasets were analyzed: (i) cruise ship outbreak-associated ANDV genome sequences generated during the international outbreak investigation^3,4^; (ii) representative Argentinian ANDV genome sequences, including previously described historic Patagonian HPS outbreak-associated lineages^13,17^; and (iii) human-derived genome sequences of ANDV collected from 2011 and 2026 from multiple endemic regions of Chile^10^. Independent datasets were generated for the ANDV S, M, and L genomic segments to enable segment-specific phylogenetic analyses.

Phylogenetic findings were interpreted together with available epidemiological and geographic information, including travel history and reported locations of probable exposure for cruise ship outbreak index cases 1 and 2. Accession numbers for all sequences included in this study are provided in **Supplementary Table S4**.

### Genome sequencing and sequence assembly

Sequencing and consensus genome sequence generation for reference genome sequences from Chile were previously described^10^. Briefly, viral RNA extracted from acute-phase human buffy coat samples was subjected to amplicon-based whole-genome amplification of the S, M, and L genomic segments following previously described orthohantavirus genome sequencing strategies^18^. For sample p1236, amplification of the second half of the L segment was technically challenging; therefore, this region was amplified using the multiplex PCR approach described by Taylor et al.^18^ and subsequently sequenced. No additional target enrichment was performed prior to sequencing.

Sequencing libraries were prepared using Nextera XT chemistry and sequenced on an Illumina iSeq 100 platform using paired-end reads (2 × 151 bp). Raw reads were quality-filtered and adapter trimmed before reference-guided assembly against the Chile-9717869 genome (GenBank: #AF291702–AF291704) using BWA-MEM. Consensus sequences were generated using iVar with a minimum base quality threshold of Q30 and minimum depth coverage of 400×. Coverage statistics were calculated using SAMtools^19^, and assembly quality was assessed using FastQC (https://www.bioinformatics.babraham.ac.uk/projects/fastqc/).

The dataset included genome sequences from Chile with high sequencing depth and near-complete genomic coverage across all three genome segments, with average genome coverage exceeding 99%. The only exception was sample p1236, for which the L segment was partially recovered, resulting in 68% coverage for that segment while retaining complete S and M genome segments.

Genome sequences associated with the cruise ship outbreak were generated independently by participating laboratories using laboratory-specific sequencing and bioinformatic workflows as previously described^3,4^.

### Phylogenetic analyses

Complete S, M, and L segment sequences were aligned independently using MAFFT v7.505^20^ and manually curated in BioEdit v7.2.5 to remove ambiguous regions and preserve functional open reading frames. Maximum-likelihood (ML) phylogenetic inference was performed in IQ-TREE v2.4^21^ using the best-fit nucleotide substitution model selected by ModelFinder^22^ under the Bayesian Information Criterion (BIC). Statistical branch support was evaluated using the SH-like approximate likelihood ratio test (SH-aLRT) with 1,000 ultrafast bootstrap replicates. Tree visualization and annotation were performed in iTOL v4^23^.

### Bayesian phylogenetic and phylogeographic analyses

Bayesian phylogenetic analyses were conducted independently for the S, M, and L genomic segments using BEAST v1.10.5^24^. Analyses implemented a codon-partitioned scheme separating codon positions 1+2 from position 3 under independent GTR+Γ substitution models with empirical base frequencies and four gamma categories. A strict molecular clock without temporal calibration was applied, such that branch lengths represented relative genetic divergence only. Tree inference was performed under a constant-size coalescent prior. Initial unconstrained analyses were conducted to evaluate the natural phylogenetic placement of outbreak-associated genome sequences. Additional analyses incorporated a monophyletic constraint for outbreak-associated sequences based on the epidemiological expectation of a single outbreak lineage. Markov chain Monte Carlo (MCMC) sampling was run for 200 million generations with enabled automatic operator optimization.

Discrete phylogeographic analyses were performed independently for the S, M, and L genome segments using BEAST v10.5.0^24^. Time-calibrated datasets were analyzed under a strict molecular clock and a constant-size coalescent tree prior. Each genome was assigned to a discrete geographic location corresponding to the most probable site of infection inferred from epidemiological investigations, including an “unknown” category when the infection site could not be confidently determined. Ancestral geographic states and migration events were reconstructed using a discrete phylogeographic framework with Markov jump counting. Maximum clade credibility trees were generated for visualization of supported geographic transitions.

Convergence and summarization were assessed using Tracer v1.7.2^25^. After discarding the first 10% of each Markov chain as burn-in, all estimated parameters achieved effective sample sizes (ESS) >200. Maximum clade credibility trees were generated using TreeAnnotator v10.5.0 and visualized in iTOL v4^23^. Well-supported geographic transitions were mapped in QGIS v3.44^26^.

Detailed model specifications, parameter settings, and convergence diagnostics are provided in the Supplementary Methods.

### Reassortment analysis

To assess concordance in evolutionary relationships across genome segments, all pairwise nucleotide p-distances were calculated independently for the S, M, and L segments among Clade III viruses. Under the null hypothesis of no reassortment, substitution counts between segment pairs were compared using a statistical framework that models the expected relationship between genetic distances while accounting for segment-specific evolutionary rates. Isolate pairs showing significant departures from the expected relationship after multiple-testing correction were further examined using sliding-window analyses and

Kolmogorov–Smirnov tests to distinguish localized from genome-wide discordance. Detailed statistical formulation, parameter estimation, filtering criteria, and significance testing are provided in the **Supplementary Methods**.

### Comparative genomic analyses

Comparative genomic analyses were performed using complete S, M, and L segment nucleotide alignments together with deduced amino acid sequences from cruise ship outbreak-associated genome sequences and representative reference genome sequences from Argentina and Chile. Segment-specific analyses included coding and non-coding regions, as well as amino acid comparisons of N, small nonstructural protein (NSs), GPC (G_N_ and G_C_), and L protein.

SNVs were identified through comparative analyses against representative reference genome sequences from Chile, including Chile-9717869, CHI-7913, and CHI-Hu13724 p1^27^. Comparative analyses additionally evaluated previously proposed candidate substitutions associated with person-to-person transmission described in Patagonia-associated HPS outbreaks^11^.

Estimates of evolutionary divergence between sequences were calculated in MEGA12 v12.0.10^28^ using the p-distance model with pairwise deletion applied to ambiguous positions. Pairwise comparisons were expressed as percentage similarity between sequences, whereas overall mean distances were used to estimate average evolutionary divergence within phylogenetically related groups.

### Structural analyses

Amino acid substitutions resulting from nonsynonymous nucleotide changes in cruise ship outbreak-associated genome sequences compared to phylogenetically related ANDV Clade III genome sequences were mapped onto available experimentally determined ANDV structures derived from the G_N_/G_C_ spikes and L protein (PDB IDs: 9P3X^29^, 2K9H^30^ and 6Q99^31^) and Alpha Fold-3 models using UCSF ChimeraX v1.11. Predicted effects of amino acid substitutions on protein stability and protein–protein interactions were assessed using DDMut and DDMut-PPI, respectively^32,33^. Structural analyses focused on the localization, solvent accessibility, domain distribution, overlap with epitopes, and alteration of *N*-glycosylation motifs.

### Geographic contextualization

Geographic contextualization analyses integrated phylogenetic findings with the spatial distribution of historical human-derived ANDV genome sequences from Chile and Argentina together with the pre-embarkation travel itinerary of the presumed index cases associated with the cruise ship outbreak. Geographic coordinates corresponding to historical cases included in the phylogenetic analyses were curated according to the most probable site of infection based on previously published epidemiological investigations^3,10^.

The pre-embarkation travel route of the presumed index case was reconstructed using available epidemiological and travel-history information collected from the Argentinian Ministry of Health Epidemiology Office and WHO during the international outbreak investigation. Mapping analyses were performed to visualize the geographic distribution of phylogenetically related genome sequences and endemic areas across southern South America.

### Ethics/data sharing

Previously determined Chilean human-derived ANDV genome sequences used as reference sequences were obtained under institutional ethics approvals as previously described^10^.

Publicly available reference sequences were retrieved from GenBank. Outbreak-associated genome sequences were facilitated by the International Genomics Consortium Investigating the *M/V Hondius* Outbreak and are publicly available at Pathoplexus.org.

## Data availability

All generated and analyzed ANDV genome sequences during this study have been deposited in GenBank under the accession numbers provided in the **Supplementary information**. Additional phylogenetic alignments, metadata, and comparative genomic datasets supporting the findings of this study are available from the corresponding authors upon reasonable request.

## Author contributions

J.H.K., G.P., N.D.T., J.A., conceived and designed the study.

H.R., C.D.-G., and P.M., led phylogenetic and phylogeographic analyses.

P.M. performed the Bayesian evolutionary analyses.

H.R., D.D.-R., R.A.-S., and N.D.T. conducted structural and comparative genomic analyses.

H.R., E.F.-L., and J.A. generated and curated genomic sequence data.

H.R., C.D.-G., D.D.-R., P.M., and N.D.T. generated figures and data visualizations.

M.F. contributed to study coordination, interpretation of epidemiological findings, and manuscript revision.

D.M.C., L.F.-B., S.P.-A., and members of the International Genomics Consortium Investigating the *M/V Hondius* Outbreak contributed genomic data, epidemiological information, and scientific expertise.

H.R., C.D.-G., J.H.K., G.P., N.D.T., and J.A. drafted the manuscript.

All authors contributed to data interpretation, critically revised the manuscript, and approved the final version.

## Additional information

**Supplementary information.** The online version contains supplementary information available at https://doi.org/xxx.

**Correspondence and requests for materials** should be addressed to P.M., G.P., N.D.T., and J.A.

***Members of the International Genomics Consortium Investigating the M/V Hondius Outbreak participating in this work***

**Argentina:** Norberto Simboli, Carlos Giovacchini (Departamento de Epidemiología, Instituto Nacional de Enfermedades Infecciosas, Administración Nacional de Laboratorios e Institutos de Salud Dr. Carlos G. Malbrán; Buenos Aires, Argentina)

**France:** Virginie Sauvage (Institut Pasteur, Université Paris Cité, Environment and Infectious Risk Research Unit - French National Reference Center for Hantaviruses), Sarah Temmam (Institut Pasteur, Université Paris Cité, Environment and Infectious Risk Research Unit - Laboratory for Urgent Response to Biological Threats (ERI-CIBU); Paris, France).

**Germany:** Maximilian Neumann (Institute for Mathematics, Heidelberg University, 69120; Heidelberg, Germany, and Institute for Biological Interfaces 5, Karlsruhe Institute of Technology, 76344 Eggenstein-Leopoldshafen; Germany).

**Netherlands:** Bas B. Oude Munnink, Marion P. G. Koopmans, David F. Nieuwenhuijse (Virology Department and Pandemic and Disaster Preparedness Research Centre, Erasmus Medical Centre; Rotterdam, Netherlands).

**Senegal:** Ibrahima Socé Fall, Moussa Moïse Diagne (Institut Pasteur de Dakar; Dakar, Senegal).

**South Africa:** Jonathan Featherston, Jacqueline Weyer (National Institute for Communicable Diseases; Sandringham, Johannesburg, South Africa).

**Spain:** Anabel Negredo (Laboratory of Arboviruses and Imported Viral Diseases., National Center of Microbiology, National Institute of Health “Carlos III”; Madrid, Spain; Ciber of Infectious Diseases, National Institute of Health “Carlos III”; Madrid, Spain), María Paz Sánchez-Seco (Laboratory of Arboviruses and Imported Viral Diseases., National Center of Microbiology, National Institute of Health “Carlos III”; Madrid, Spain).

**Switzerland:** Florian Laubscher, Samuel Cordey (Laboratory of Virology, Laboratory Medicine Division, Diagnostic Department, Geneva University Hospitals; Geneva, Switzerland), Lorenzo Subissi, Olivier Le Polain de Waroux, Boris I. Pavlin, Ana Hoxha (WHO Health Emergencies Programme; Geneva, Switzerland).

**United States:** Qingyuan Cai (Program of Mathematical Genomics, Department of Systems Biology, Columbia University, New York, NY, USA), Raul Rabadan (Program of Mathematical Genomics, Department of Systems Biology, Columbia University, and Department of Biomedical Informatics, Columbia University, New York, NY, USA).

