## Supplementary Information for "A historical cross-border Andes virus lineage reveals the origin of a cruise ship hantavirus pulmonary syndrome outbreak"

**for**

### Supplementary Figures

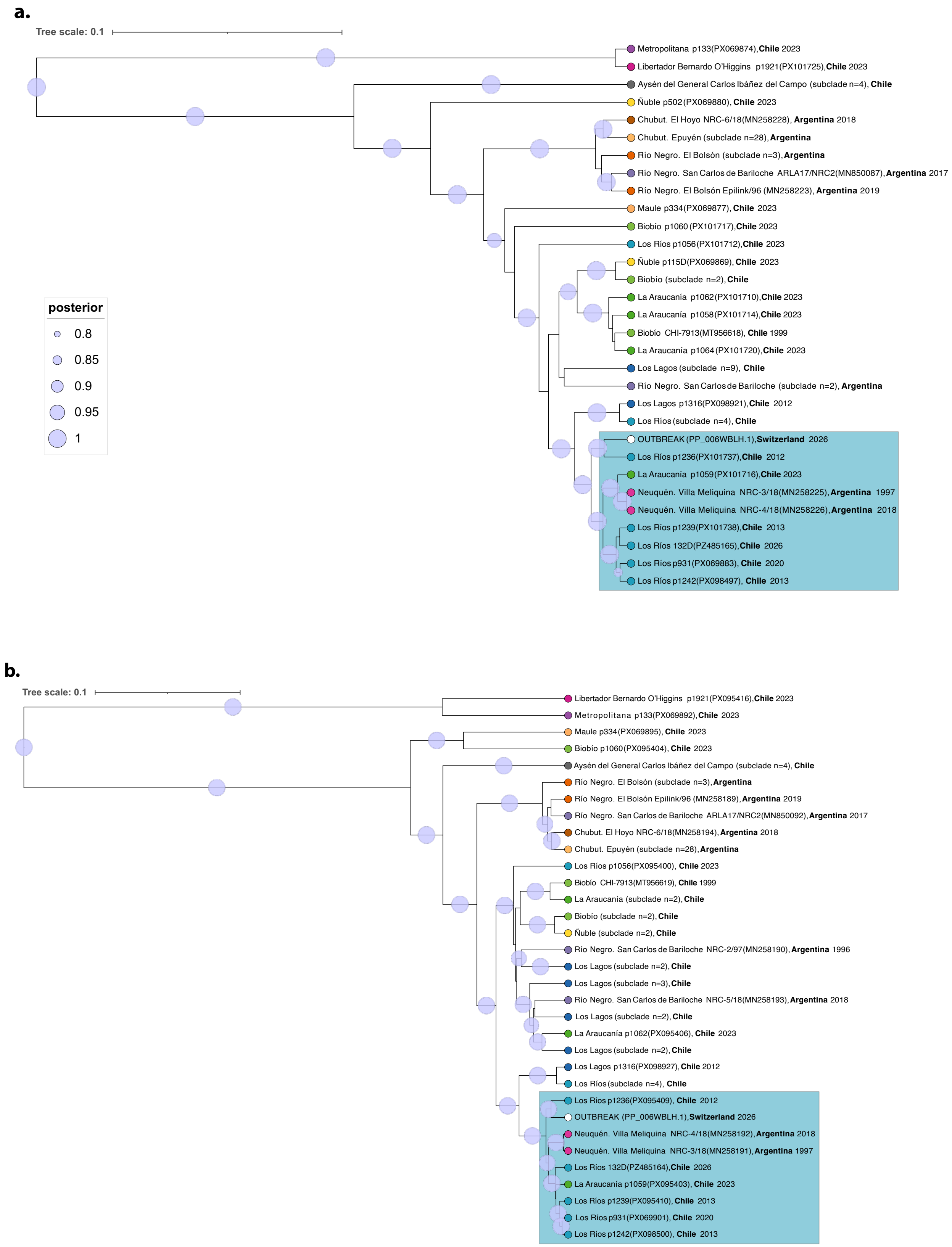

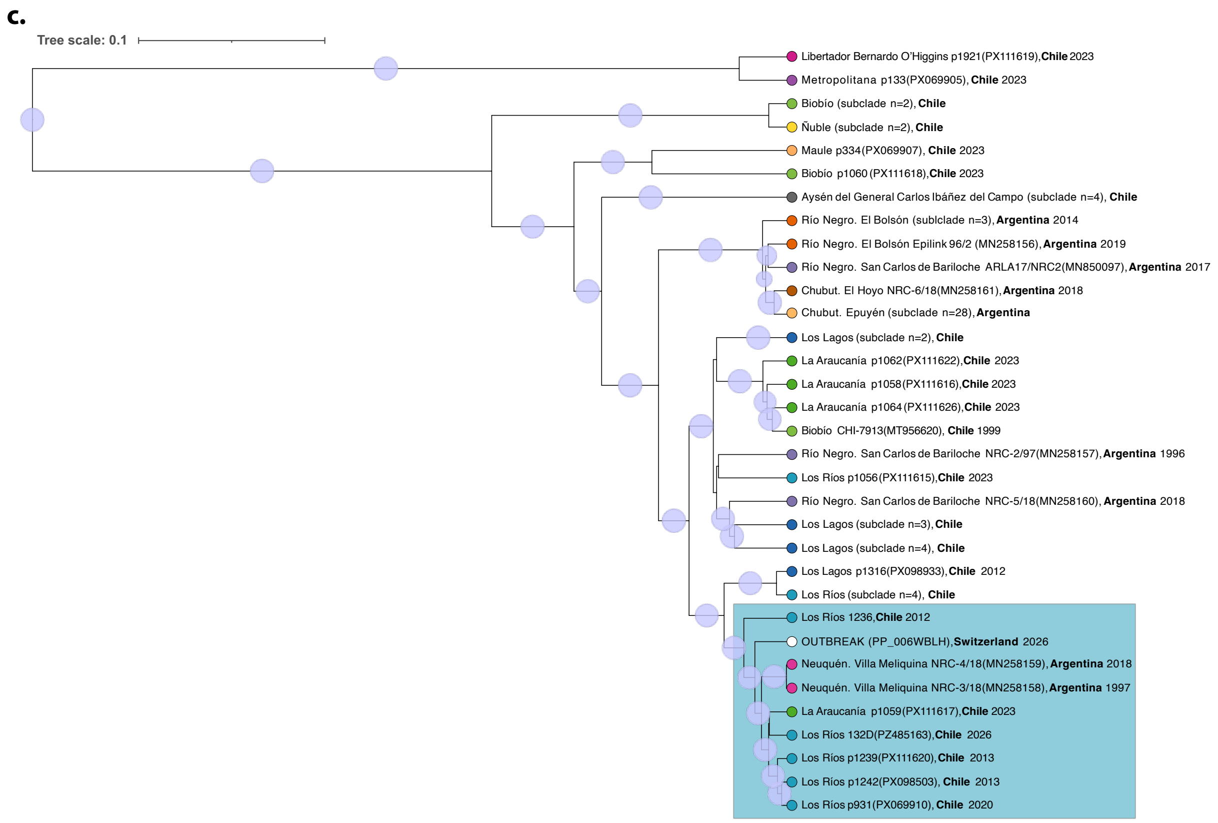


***Supplementary Fig. S1 | Cruise ship outbreak-associated Andes virus genome sequences cluster with those of Andes viruses circulating in the Hua Hum Pass corridor: Bayesian maximum clade credibility phylogenies of complete Andes virus genomes associated with the 2026 cruise ship outbreak.*** Maximum clade credibility (MCC) trees were inferred independently from Bayesian phylogenetic analyses of complete (**a**) S, (**b**) M, and (**c**) L genomic segment sequences of ANDV, including a single representative ANDV genome sequence from the cruise ship outbreak together with complete ANDV genome sequences derived from rodents, human cases, and cultured viral isolates from Argentina and Chile[10]. Sequence labels are color-coded according to geographic origin. Node colors represent posterior probability support according to the scale shown in the legend.

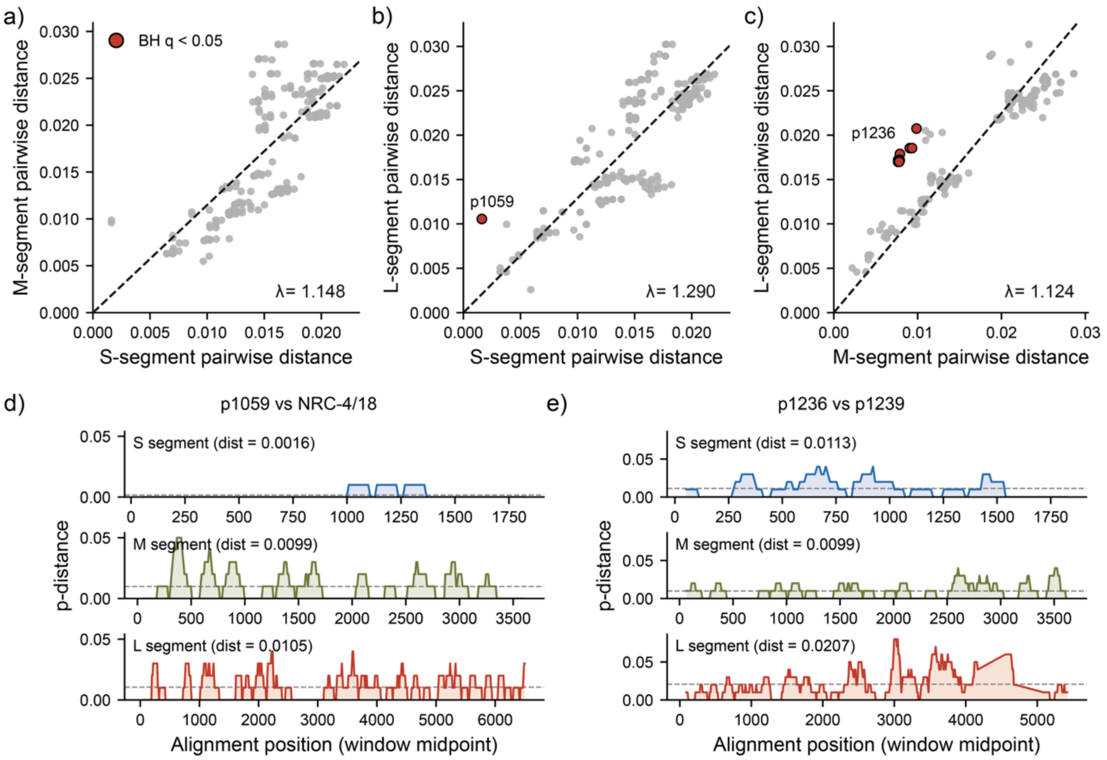


***Supplementary Fig. S2 | Cruise ship outbreak-associated Andes virus genome sequences cluster with those of Andes viruses circulating in the Hua Hum Pass corridor: P-distance analysis hardly detect evidence of reassortment among Clade III genomes.*** (a–c) Pairwise p-distances between Clade III viruses for (a) S vs M, (b) S vs L and (c) M vs L. The dashed line is $y = \lambda x$, in which $\lambda$ is the ML mutation-rate ratio between the indicated segments estimated from all pairs with sufficient variations. Points outlined in black deviate from this expectation at Benjamini–Hochberg $q$ <0.05. Significant outliers involve p1059 and/or p1236 and are examined by sliding-window analysis in (d,e). (d,e) Sliding-window pairwise p-distances (100-nt window, 10-nt stride) for (d) p1059 vs NRC-4/18 and (e) p1236 vs p1239. Dashed lines indicate overall segment p-distances. Both comparisons show some artifacts that could explain the deviation, where in (d), L-segment distance is elevated only in the 3′ half; and in (e), the S segment of p1059 is nearly identical to NRC-4/18.

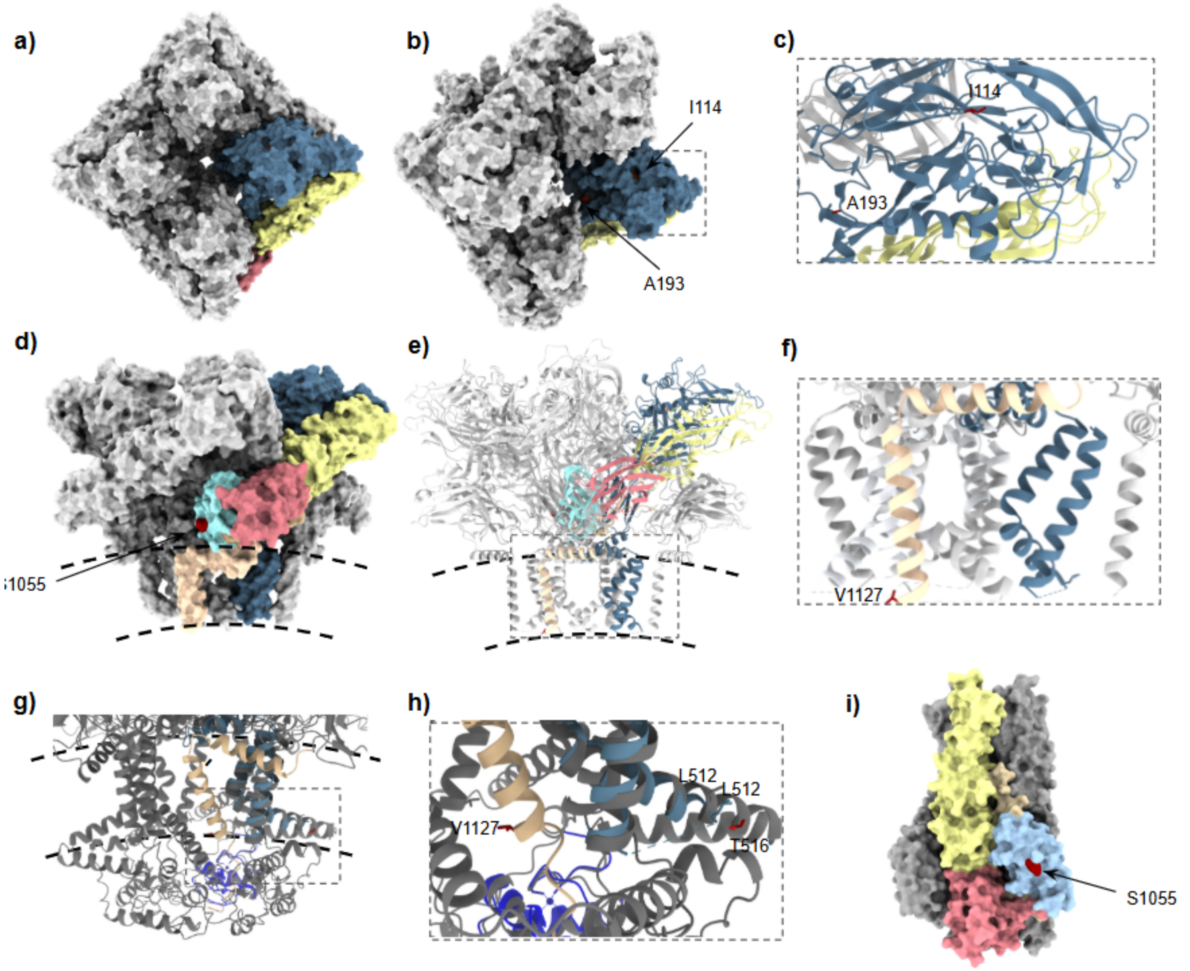


***Supplementary Fig. S3 | The Hua Hum Pass corridor Andes virus cluster is defined by limited, largely neutral molecular signatures within Andes virus Clade III diversity.*** (**a–b**) Top view of the Andes virus (ANDV) glycoprotein $G_N$/$G_C$ subunit tetramer in surface representation (PDB ID: 9P3X)[1]. A single heterodimer is highlighted by domains: the $G_N$ monomer is shown in slate blue, and the $G_C$ monomer is color-coded by domains (D): DI in red, DII in yellow, DIII in light blue, and the stem, pre-transmembrane region and transmembrane domain (TMD) in beige. The remaining three heterodimers are rendered in light grey. The specific residue positions I114 and A193, located on the surface of the $G_N$ head, are highlighted in red. (**c**) Close-up of from (b) of the $G_N$ region showing the detailed orientation of residues I114 and A193 in cartoon representation. (**d–e**) Side view of the $G_N$/$G_C$ tetramer in surface (d) and cartoon representation (**e**), highlighting the residue S1055 on DIII and V1127 in the TMD in stick representation in red. (**f**) Close-up view of the TMDs from (e) showing V1127 in stick representation at the resolved C-terminal end of the

$G_C$ TMD. (**g**) Structural alignment of the transmembrane region and endodomain from an AlphaFold 3-predictive model (grey)[33] superimposed onto the $G_N$ TMDs of the cryo-EM structure of the 9P3X heterodimer ($G_N$ TMDs in slate blue, $G_C$ TMD in beige) and the structure of the homologous $G_N$ zinc-finger domain (PDB ID: 2K9H)[2] in dark blue. (**h**) Close-up view of the aligned region highlighting the structural environment of residues V1127 and T516 in red in stick representation; the spatial convergence of the conserved L512 residues from both the predictive model and the experimental structure is also shown in grey and slate blue stick representation, respectively. (**i**) Side view of the $G_C$ subunit in its post-fusion trimeric state conformation, shown in surface representation (PDB ID: 6Y6Q)[3]. Color code as (a). The surface accessibility in the $G_C$ post-fusion trimer of residue S1055 on DIII is indicated in red.

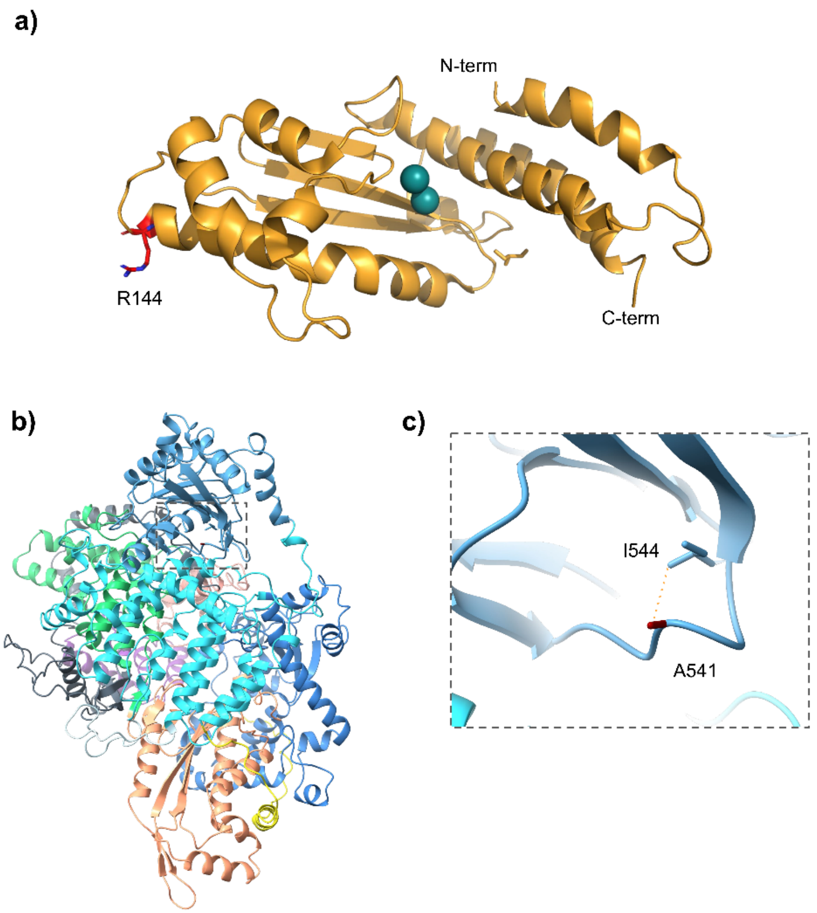


***Supplementary Fig. S4 | The Hua Hum Pass corridor Andes virus cluster is defined by limited, largely neutral molecular signatures within Andes virus Clade III diversity.*** (**a**) N-terminal endonuclease domain (aa 1–200) (PDB ID: 6Q99)[4] in cartoon representation displays the positively charged R144K substitution rendered as a stick in red. The manganese ions are bound to the active site and represented as green spheres. (**b**) Alpha Fold-3 model of the ANDV L protein core region (aa 226–1,600) based on SNV L protein (PDB: 8CI5)[5] depicted as a cartoon diagram. The structural domains and subdomains are color-coded based on their primary sequence intervals as reported previously[5]: the linker region (aa 226–263, yellow); the discontinuous core lobe domain (aa 264–292, 391–471, 565–675, and 700–735, turquoise); the interspersed viral RNA-binding lobe (vRBL) subdomains (aa 293–390 and 472–564, steel blue); the discontinuous fingers (aa 736–920 and 983–1,058, royal blue); the structural palm (aa 921–982 and 1,059–1,181, light salmon); the thumb (aa 1,182–1,341, green); the extended bridge region (aa 1,342–1,400, salmon); the alternating thumb ring subdomains (aa 1,401–1,493 and 1,565–1,600, grey); and the intervening lid region (aa 1,503–1,565, purple). (**c**) Close-up view from (b) of the viral RNA binding lobe showing the location of substitution A541V, displayed as stick in red. The hydrophobic contact established between the A541 side chain and the neighboring I544 residue is shown as a dashed yellow line.

#### Supplementary Tables

***Supplementary Table S1 | Cruise ship outbreak-associated Andes virus genome sequences cluster with those of Andes viruses circulating in the Hua Hum Pass corridor: Epidemiological and clinical characteristics of Chilean human-derived Andes virus cases belonging to the Hua Hum Pass corridor cluster***

| Year of sample collection | Case ID | Locality | Exposure setting | Epidemiological context | HPS severity[a] |
|---|---|---|---|---|---|
| 2012 | p1236 | Los Ríos Region (Lanco) | Rural exposure | Isolated case | Severe |
| 2013 | p1239 | Los Ríos Region (San José de Mariquina) | Rural exposure | Isolated case | Mild |
| 2013 | p1242 | Los Ríos Region (Lanco) | Rural exposure | Isolated case | Mild |
| 2020 | p931 | Los Ríos Region (Valdivia) | Environmental household exposure | Family-associated cluster | Mild |
| 2023 | p1059 | Araucanía Region (Curarrehue) | Rural exposure | Isolated case | Mild |
| 2026 | p132D | Los Ríos Region (Licanray) | Rural exposure | Isolated case | Mild |

[a]Severity classification followed previously described nationwide Chilean hantavirus pulmonary syndrome (HPS) criteria: mild disease includes patients requiring supplemental oxygen without invasive mechanical ventilation, whereas severe disease includes cases requiring vasoactive support, invasive mechanical ventilation, or extracorporeal membrane oxygenation (ECMO)[6,7].

***Supplementary Table S2 | The Hua Hum Pass corridor Andes virus cluster is defined by limited, largely neutral molecular signatures within Andes virus Clade III diversity: Deduced amino acid signatures distinguishing Hua Hum Pass corridor cluster Andes virus within Clade III Andes virus diversity***

| **Substitution** | **HHPC cluster[a] (*n*=9)** | **non-HHPC cluster Clade III[b] (*n*=9)** | **Remaining dataset (*n*= 64)** | **Classification** |
|---|---|---|---|---|
| **S segment** | | | | |
| **N protein (aa position)[c]** | | | | |
| **L7I** | Yes (5/9) | No | No | **HHPC cluster-enriched** |
| M segment | | | | |
| GPC ($G_N$/$G_c$ subunits) (aa position)[c] | | | | |
| **I114V** | Yes (9/9) | Yes (9/9) | Yes (1/64) | Clade III-enriched |
| **A193T** | Yes (2/9) (NRC-3/18, NRC-4/18) | No | No | Rare HHPC cluster variant |
| **T516I** | Yes (1/9) (All cruise ship sequences) | No | No | **Outbreak-specific** |
| **S1055T** | Yes (9/9) | No | No | **HHPC cluster-defining marker** |
| **S1055A** | No | Yes (9/9) | No | Non-HHPC cluster Clade III-defining marker |
| **V1127I** | Yes (9/9) | Yes (9/9) | Yes (4/64) | Clade III-enriched |
| **L segment** | | | | |
| **L protein (aa position)[c]** | | | | |

| | | | | |
|---|---|---|---|---|
| **I70V** | Yes (2/9) | No | Yes (2/64) *ANDV Chi-North | Not HHPC cluster-specific |
| **R144K** | Yes (7/9) (Not in cruise ship sequences) | Yes (9/9) | Yes (4/64) *2 ANDV Chi-North | Clade III-enriched |
| **P149S** | Yes (3/9) (Not in cruise ship sequences) | No | Yes (1/64) *ANDV Chi-North | Not HHPC cluster-specific |
| **A541V** | Yes (9/9) | Yes (9/9) | Yes (13/64) | Clade III-enriched |

[a]**Hua Hum Pass corridor (HHPC) cluster:** p132D, p931, p1059, p1236, p1239, p1242, NRC-3/18, NRC-4/18, and cruise ship outbreak sequences (*n*=10, represented as a single consensus sequence considered as 1).

[b]**Clade III:** HHPC cluster sequences plus those of p136, p137, p1316, CHI-Hu13724 (P1 and P2), ARG-M.a., ARG p3, ARG p9, and ARG p19.

[c]**Inclusion criteria**: single nucleotide variants (SNVs) in at least two sequences of the HHPC cluster. The analysis included a consensus sequence derived from the ten outbreak-associated genome sequences. Individual examination of each genome sequence did not reveal additional non-synonymous SNVs among the cruise ship outbreak sequences. Colored cells highlight the two principal HHPC cluster-associated markers (L7I and S1055T) and the cruise ship outbreak-specific marker (T516I).

***Supplementary Table S3 | The Hua Hum Pass corridor cluster is defined by limited, largely neutral molecular signatures within Andes virus Clade III diversity: Structural mapping of single nucleotide variants identified within Hua Hum Pass corridor cluster.*** Comparative analysis of deduced amino acid substitutions identified among the Hua Hum Pass corridor cluster genome sequences, representative Andes virus clades (I–V), and selected orthohantavirus genome sequences.

| | **ANDV** | | | | | | | | | | | | | | **Other orthohantaviruses** | | | |
|---|---|---|---|---|---|---|---|---|---|---|---|---|---|---|---|---|---|---|
| | **P133 (ANDV Chi-North)** | **CHI-9717869** | **Strain Epuyén** | **CHI-7913** | **CHI-Hu13724** | **p132D** | **p931** | **p1059** | **p1236** | **p1239** | **p1242** | **NRC-3/18** | **NRC-4/18** | **Cruise ship outbreak** | **ORNV (22996)** | **SNV (NM H10)** | **HTNV (HV004)** | **Domain/Function** |
| **Sample characteristics** | | | | | | | | | | | | | | | | | | |
| **Country** | Chile | Chile | Argentine | Chile | Chile | Chile | Chile | Chile | Chile | Chile | Chile | Argentina | Argentina | | Argentina | United States | China | |
| **Geographic origin** | Metropolitana | Coyhaique, Aysén | Epuyén | Mulchén, BioBío | Los Ríos | Los Ríos | Los Ríos | La Araucanía | Los Ríos | Los Ríos | Los Ríos | Neuquén | Neuquén | | Orán, Salta | New Mexico | Hubei | |
| **Clade** | V | IV | II | I | III | III | III | III | III | III | III | III | III | III | ANDV-like | - | - | |
| **Genomic region (aa position)** | | | | | | | | | | | | | | | | | | |
| **S segment** | | | | | | | | | | | | | | | | | | |
| **GenBank #** | PX069874 | AF291702 | PQ215668 | MT956618 | PV808473 | PZ485165 | PX069883 | PX101716 | PX101737 | PX101738 | PX098497 | MN258225 | MN258226 | | AF482715 | NC_005216 | JQ083395 | |
| **N protein** | | | | | | | | | | | | | | | | | | |
| L7I | L | L | L | L | L | I | I | L | L | I | I | L | L | L | L | V | L | N-terminal coiled-coil |

| | | | | | | | | | | | | | | | | | |
|---|---|---|---|---|---|---|---|---|---|---|---|---|---|---|---|---|---|
| | | | | | | | | | | | | | | | | | (RdRp binding*) |
| **NSs protein** | | | | | | | | | | | | | | | | | |
| Q5R | R | Q | R | R | R | R | R | R | R | R | R | R | R | R | R | R | N/A | |
| E33G | E | E | G | G | G | G | G | G | G | G | G | G | G | G | E | E | N/A | |
| L35S | S | L | S | S | L | S | S | S | S | S | S | S | S | S | L | L | N/A | |
| G37D | G | G | G | D | G | G | G | G | G | G | G | G | G | G | G | G | N/A | |
| Q40R | R | Q | R | Q | Q | Q | Q | Q | Q | Q | Q | Q | Q | Q | R | L | N/A | |
| N47S | N | N | S | N | N | N | N | N | N | N | N | N | N | N | N | N | N/A | |
| I62T | I | I | T | T | T | T | T | T | T | T | T | T | T | T | I | T | N/A | |
| **M segment** | | | | | | | | | | | | | | | | | |
| **GenBank #** | PX069892 | AF291703 | PQ215669 | MT956619 | PV808474 | PZ485164 | PX069901 | PX095403 | PX095409 | PX095410 | PX098500 | MN258159 | MN258192 | | AF028024 | NC_005215 | JQ083394 | |
| **GPC** | | | | | | | | | | | | | | | | | |
| V8A | A | V | A | A | A | A | A | A | A | A | A | A | A | A | V | F | V | $G_N$ signal peptide (semi-conserved) |
| T14M | T | T | T | T | T | T | T | T | T | T | T | T | T | T | T | T | W | $G_N$ signal peptide (semi-conserved) |
| **I114V** | **I** | **I** | **I** | **I** | **V** | **V** | **V** | **V** | **V** | **V** | **V** | **V** | **V** | **V** | **I** | **N** | **K** | **$G_N$ head (partially e, hypervariable)** |

| | | | | | | | | | | | | | | | | | | |
|---|---|---|---|---|---|---|---|---|---|---|---|---|---|---|---|---|---|---|
| **A193T** | **A** | **A** | **A** | **A** | **A** | **A** | **A** | **A** | **A** | **A** | **A** | **T** | **T** | **A** | **S** | **T** | **V** | **$G_N$ head (exposed, variable)** |
| F216L | F | F | L | F | F | F | F | F | F | F | F | F | F | F | F | L | K | $G_N$ head (b, variable) |
| K236R | K | K | K | K | K | K | R | K | K | K | K | K | K | K | K | G | D | $G_N$ head (exposed, variable) |
| R281K | R | R | R | R | R | R | R | R | **K** | R | R | R | R | R | R | R | D | $G_N$ Head, (b neutral) IA S236 (3.3 Å) |
| H294Y | Y | H | Y | Y | Y | Y | Y | Y | Y | Y | Y | Y | Y | Y | H | L | I | $G_N$ head (variable), $G_C$ interface, IA T734 (3.1 Å) |
| V346I | I | V | I | I | I | I | I | I | I | I | I | I | I | I | L | I | L | $G_N$ head, (b, semi conserved) at $G_C$ interface |
| T353V | V | T | I | V | V | V | V | V | V | V | V | V | V | V | V | V | K | $G_N$ head (e, hypervariable) |
| V435I | V | V | V | V | V | I | V | V | V | V | V | V | V | V | V | V | C | $G_N$ base (b, variable) |
| V499I | I | V | I | V | V | V | V | V | V | V | V | V | V | V | I | L | p | $G_N$ transmembrane 1 (b, semi conserved) |
| **T516I** | **T** | **T** | **T** | **T** | **T** | **T** | **T** | **T** | **T** | **T** | **T** | **T** | **T** | **I** | **T** | **T** | **F** | **Endodomain (predicted at membrane interface, semi conserved)** |
| I537V | V | I | V | V | V | V | V | V | V | V | V | V | V | V | V | V | E | Endodomain (b, variable) |

| | | | | | | | | | | | | | | | | | | |
|---|---|---|---|---|---|---|---|---|---|---|---|---|---|---|---|---|---|---|
| K614R | K | K | K | K | K | K | K | K | K | K | R | K | K | K | K | R | C | Endodomain (b, variable) |
| T641I | T | T | I | T | T | T | T | T | T | T | T | T | T | T | T | T | S | $G_N$ transmembrane 2 (b, variable) |
| E933D | E | E | E | E | E | E | E | E | E | E | E | E | E | E | D | E | H | $G_C$ ectodomain, on DII surface (b, $G_N$2 interface, variable) |
| T938A | T | T | A | A | T | T | T | T | T | T | T | T | T | T | T | S | R | $G_C$ ectodomain, on DII loop (e, $G_N$2 interface, hypervariable) |
| T994A | T | T | T | T | T | T | T | T | T | T | T | T | A | T | T | V | L | $G_C$ ectodomain, on DIII surface (b in prefusion, semiconserved) |
| T1023A | A | T | A | A | A | A | A | A | A | A | A | A | A | A | T | T | L | $G_C$ ectodomain, DIII (buried in perfusion, on DIII surface, neutral) |
| **S1055T** | **S** | **S** | **S** | **S** | **A** | **T** | **T** | **T** | **T** | **T** | **T** | **T** | **T** | **T** | **S** | **E** | **G** | **$G_C$ ectodomain, DIII (on DII surface, b in prefusion, hypervariable)** |

| | | | | | | | | | | | | | | | | | | |
|---|---|---|---|---|---|---|---|---|---|---|---|---|---|---|---|---|---|---|
| V1115I | V | V | I | V | V | V | V | V | V | V | V | V | V | V | I | I | L | $G_c$ transmembrane (b, variable) |
| **V1127I** | **I** | **V** | **V** | **V** | **I** | **I** | **I** | **I** | **I** | **I** | **I** | **I** | **I** | **I** | **I** | **F** | **C** | **$G_c$ transmembrane (b, neutral)** |
| **L segment** | | | | | | | | | | | | | | | | | | |
| **GenBank #** | PX069905 | AF291704 | PQ215670 | MT956620 | PV808475 | PZ485163 | PX069910 | PX111617 | PZ568904 | PX111620 | PX098503 | MN258158 | MN258159 | | - | NC_005217 | JQ083393 | |
| **RdRp** | | | | | | | | | | | | | | | | | | |
| I70V | V | I | I | I | I | I | I | I | I | I | I | V | V | I | | V | I | Endonuclease, cap-snatching (semiconserved) |
| T141I | I | T | V | I | I | I | I | I | I | I | I | I | I | I | | Q | K | Endonuclease, cap-snatching (hypervariable) |
| R144K | K | R | R | R | K | R | K | K | K | K | K | K | K | R | | K | L | Endonuclease, cap-snatching (hypervariable) |
| P149S | P | P | P | P | P | P | S | P | P | S | S | P | P | P | | N | P | Endonuclease, cap-snatching (hypervariable) |
| S273N | N | S | S | S | S | N | S | S | S | S | S | S | S | S | | N | S | Core lobe (hypervariable) |
| Q276R | Q | Q | Q | Q | Q | R | Q | Q | Q | Q | Q | Q | Q | Q | | S | A | Core lobe (hypervariable) |

| | | | | | | | | | | | | | | | | | | |
|---|---|---|---|---|---|---|---|---|---|---|---|---|---|---|---|---|---|---|
| S277L | V | L | S | S | S | S | S | S | S | S | S | S | S | S | | V | N | Core lobe (hypervariable) |
| Q280E | Q | Q | Q | Q | Q | E | Q | Q | Q | Q | Q | Q | Q | Q | | S | Q | Core lobe (neutral) |
| K281R | K | K | K | K | K | K | K | K | K | K | K | K | K | K | | R | K | Core lobe (hypervariable) |
| A338S | A | S | A | A | A | A | A | A | A | A | A | A | A | A | | A | A | Core lobe, vRBL (semiconserved) |
| K346R | K | R | K | K | K | K | K | K | K | K | K | K | K | K | | R | R | Core lobe, vRBL (variable) |
| N364S | N | N | S | N | N | N | N | N | N | N | N | N | N | N | | S | N | Core lobe, vRBL (variable) |
| V402I | V | I | I | V | V | V | V | V | V | V | V | V | V | V | | P | I | Core lobe, vRBL (hypervariable) |
| A541V | A | A | A | A | V | V | V | V | V | V | V | V | V | V | | E | K | Core lobe, vRBL (hypervariable) |
| A581S | A | A | A | A | S | A | A | A | A | A | A | A | A | A | | A | A | Core lobe (conserved) |
| V614M | M | V | V | V | V | M | V | V | V | V | V | V | V | V | | V | C | Core lobe (neutral) |
| N780D | E | D | N | N | N | N | N | N | D | N | N | N | N | N | | Q | E | Core lobe, fingers (hypervariable) |

| | | | | | | | | | | | | | | | | | | |
|---|---|---|---|---|---|---|---|---|---|---|---|---|---|---|---|---|---|---|
| S876A | S | S | A | S | S | S | S | S | S | S | S | S | S | S | | H | K | Core lobe, fingers (variable) |
| D1033N | E | N | D | D | D | D | D | D | D | D | D | D | D | D | | E | S | RdRp core, fingers (hypervariable ) |
| V1073I | I | V | V | V | V | I | V | V | V | V | V | V | V | V | | V | S | RdRp core, palm (neutral) |
| K1082R | K | K | K | K | K | K | K | K | K | K | K | K | K | K | | A | N | RdRp core, palm (hypervariable ) |
| N1288I | N | N | N | N | N | N | N | N | I | N | N | N | N | N | | N | L | RdRp core, thumb (neutral) |
| E1303D | D | D | E | E | E | E | E | E | E | E | E | E | E | E | | D | K | RdRp core, thumb (hypervariable ) |
| Y1420F | Y | Y | Y | Y | Y | Y | Y | Y | Y | F | Y | Y | Y | Y | | L | L | RdRp core, thumb ring (hypervariable ) |
| S1440N | S | S | N | S | S | S | S | S | S | S | S | S | S | S | | K | Y | RdRp core, thumb ring (hypervariable ) |
| I1665V | I | V | V | I | I | I | I | I | V | I | I | I | I | I | | V | L | C-terminal (variable) |
| R1750K | K | K | R | R | R | R | R | R | ND[a] | R | R | R | R | R | | K | R | C-terminal (cap-binding) (hypervariable ) |

| | | | | | | | | | | | | | | | | | | |
|---|---|---|---|---|---|---|---|---|---|---|---|---|---|---|---|---|---|---|
| R1939K | R | R | R | R | R | R | R | R | ND[a] | K | R | R | R | R | | P | P | C-terminal (semiconserved) |
| N1940K | N | N | N | N | N | N | N | N | ND[a] | N | N | N | N | N | | R | N | C-terminal (hypervariable) |
| Q1965H | Q | Q | H | Q | Q | Q | Q | Q | ND[a] | Q | Q | Q | Q | Q | | Q | W | C-terminal (variable) |
| K2027R | K | K | K | K | K | K | K | K | ND[a] | K | K | K | K | K | | R | R | C-terminal (neutral) |
| D2106G | E | D | D | D | D | D | D | D | ND[a] | D | G | D | D | D | | E | S | C-terminal (variable) |
| I2109V | M | I | V | V | I | I | I | I | ND[a] | I | I | I | I | I | | N | R | C-terminal (hypervariable) |
| A2113T | T | T | T | A | A | A | A | A | ND[a] | A | A | A | A | A | | R | N | C-terminal (hypervariable) |
| K2116E | K | K | E | K | K | K | K | K | ND[a] | K | K | K | K | K | | E | E | C-terminal (hypervariable) |

Representative isolates from the four majors Chilean ANDV clades (Clade I: CHI-7913; Clade II: Strain Epuyén; Clade III: CHI-Hu13724; Clade IV: Chile-9717869) and representative Orán, Sin Nombre, and Hantaan viruses were included for comparison. The representative sequence for Clade V (ANDV-CHI-North) was selected based on its highest similarity to the consensus sequence of the Clade V genomes included in this study and was confirmed to be representative of the complete Clade V dataset[10]. The cruise ship outbreak column includes the seven available viral genomes across all segments, as they share identical substitution patterns: (PP_006W3U9.2, PP_006W6RC.2, PP_006WBLH.2, PP_006WDJK.1, PP_006WDKH.1, PP_006XBKH.1, PP_006XDJH.5, PP_006XDHK.5, PP_0702S6, PP_0072Q1X). Only amino acid substitutions identified within the ANDV-south dataset were considered, excluding differences unique to ANDV-Chi-North and non-ANDV orthohantaviruses; all substitutions detected in at least one ANDV sequence are shown. The Domain/Function column indicates known structural or functional annotations when available. Conservation categories for N, GPC, and RdRp were inferred using Consurf, ranging from hypervariable to conserved. For GPC, residue accessibility is reported as exposed (e) or buried (b) based on Consurf predictions and manual inspection of available experimental structures. Substitutions specific to the cruise ship outbreak are highlighted in bold. [a] ND = amino acid could not be determined because of insufficient sequencing coverage.

***Supplementary Table S4 | Study overview and datasets: GenBank accession numbers of all genome sequences included in this study***

| Sequence ID | S segment | M segment | L segment |
|---|---|---|---|
| 127D | PZ485159 | PZ485158 | PZ485159 |
| 131D | PZ485162 | PZ485161 | PZ485162 |
| 132D | PZ485165 | PZ485164 | PZ485165 |
| 136D | PZ485168 | PZ485167 | PZ485168 |
| ANDV ARCH14/NRC1 | MN850086.1 | MN850091.1 | MN850096.1 |
| ANDV AREB14/P1 | MN850083.1 | MN850088.1 | MN850093.1 |
| ANDV AREB14/P2 | MN850084.1 | MN850089.1 | MN850094.1 |
| ANDV AREB14/P3 | MN850085.1 | MN850090.1 | MN850095.1 |
| ANDV ARG p19 | OP555722.1 | OP555727.1 | OP555734.1 |
| ANDV ARG p3 | OP555720.1 | OP555725.1 | OP555732.1 |
| ANDV ARG p9 | OP555721.1 | OP555726.1 | OP555733.1 |
| ANDV ARG-M.a. | OP555728.1 | OP555729.1 | OP555735.1 |
| ANDV ARLA17/NRC2 | MN850087.1 | MN850092.1 | MN850097.1 |
| ANDV CHI-7913 | MT956618.1 | MT956619.1 | MT956620.1 |
| ANDV CHI-Hu13724 P1 | PV808473.1 | PV808474.1 | PV808475.1 |
| ANDV CHI-Hu13724 P2 | PV808476.1 | PV808477.1 | PV808478.1 |
| ANDV Chile-9717869 | MT956622.1 | MT956623.1 | MT956621.1 |
| ANDV Epilink/96 | MN258223.1 | MN258189.1 | MN258156.1 |
| ANDV Epuyén | PQ215668.1 | PQ215669.1 | PQ215670.1 |
| ANDV Epuyén/18–19_Patient_1 | MN258239.1 | MN258205.1 | MN258172.1 |
| ANDV Epuyén/18–19_Patient_10 | MN258229.1 | MN258195.1 | MN258162.1 |
| ANDV Epuyén/18–19_Patient_11 | MN258230.1 | MN258196.1 | MN258163.1 |
| ANDV Epuyén/18–19_Patient_12 | MN258231.1 | MN258197.1 | MN258164.1 |
| ANDV Epuyén/18–19_Patient_13 | MN258232.1 | MN258198.1 | MN258165.1 |
| ANDV Epuyén/18–19_Patient_14 | MN258233.1 | MN258199.1 | MN258166.1 |
| ANDV Epuyén/18–19_Patient_15 | MN258234.1 | MN258200.1 | MN258167.1 |
| ANDV Epuyén/18–19_Patient_16 | MN258235.1 | MN258201.1 | MN258168.1 |
| ANDV Epuyén/18–19_Patient_17 | MN258236.1 | MN258202.1 | MN258169.1 |
| ANDV Epuyén/18–19_Patient_18 | MN258237.1 | MN258203.1 | MN258170.1 |
| ANDV Epuyén/18–19_Patient_19 | MN258238.1 | MN258204.1 | MN258171.1 |
| ANDV Epuyén/18–19_Patient_20 | MN258240.1 | MN258206.1 | MN258173.1 |
| ANDV Epuyén/18–19_Patient_22 | MN258241.1 | MN258207.1 | MN258174.1 |
| ANDV Epuyén/18–19_Patient_23 | MN258242.1 | MN258208.1 | MN258175.1 |
| ANDV Epuyén/18–19_Patient_24 | MN258243.1 | MN258209.1 | MN258176.1 |
| ANDV Epuyén/18–19_Patient_25 | MN258244.1 | MN258210.1 | MN258177.1 |
| ANDV Epuyén/18–19_Patient_26 | MN258245.1 | MN258211.1 | MN258178.1 |
| ANDV Epuyén/18–19_Patient_27 | MN258246.1 | MN258212.1 | MN258179.1 |

| | | | |
|---|---|---|---|
| ANDV Epuyén/18–19_Patient_28 | MN258247.1 | MN258213.1 | MN258180.1 |
| ANDV Epuyén/18–19_Patient_29 | MN258248.1 | MN258214.1 | MN258181.1 |
| ANDV Epuyén/18–19_Patient_3 | MN258250.1 | MN258216.1 | MN258182.1 |
| ANDV Epuyén/18–19_Patient_4 | MN258251.1 | MN258217.1 | MN258183.1 |
| ANDV Epuyén/18–19_Patient_5 | MN258252.1 | MN258218.1 | MN258184.1 |
| ANDV Epuyén/18–19_Patient_6 | MN258253.1 | MN258219.1 | MN258185.1 |
| ANDV Epuyén/18–19_Patient_7 | MN258254.1 | MN258220.1 | MN258186.1 |
| ANDV Epuyén/18–19_Patient_8 | MN258255.1 | MN258221.1 | MN258187.1 |
| ANDV Epuyén/18–19_Patient_9 | MN258256.1 | MN258222.1 | MN258188.1 |
| ANDV NRC-2/97 | MN258224.1 | MN258190.1 | MN258157.1 |
| ANDV NRC-3/18 | MN258225.1 | MN258191.1 | MN258158.1 |
| ANDV NRC-4/18 | MN258226.1 | MN258192.1 | MN258159.1 |
| ANDV NRC-5/18 | MN258227.1 | MN258193.1 | MN258160.1 |
| ANDV NRC-6/18 | MN258228.1 | MN258194.1 | MN258161.1 |
| ANDV-CH-LS-2016 | OR405525.1 | OR405524.1 | OR405523.1 |
| ANDV-CH-LS-2022 | OR405520.1 | OR405521.1 | OR405522.1 |
| Canada-Case 11 blood | PP 006XDHK.6 | PP 006XDJH.6 | PP 006XDHK.6 |
| Canada-Case 11 NPswab | PP 006XDJH.6 | PP 006XDHK.6 | PP 006XDJH.6 |
| France-Case 9 | PP 006XBKH.2 | PP 006XBKH.2 | PP 006XBKH.2 |
| Johannesburg-Case 2 | PP 006WDKH.3 | PP 006WDKH.3 | PP 006WDKH.3 |
| Johannesburg-Case 3 | PP 006WDJK.3 | PP 006WDJK.3 | PP 006WDJK.3 |
| Netherlands-Case 5 | PP 006W3U9.2 | PP 006W6RC.3 | PP 006W3U9.2 |
| Netherlands-Case 6 | PP 006W6RC.3 | PP 006W3U9.2 | PP 006W6RC.3 |
| p1052 | PX101710 | PX095398 | PX069911 |
| p1056 | PX101712 | PX095400 | PX111615 |
| p1058 | PX101714 | PX095402 | PX111616 |
| p1059 | PX101716 | PX095403 | PX111617 |
| p1060 | PX101717 | PX095404 | PX111618 |
| p1062 | PX101719 | PX095406 | PX111622 |
| p1064 | PX101720 | PX095407 | PX111626 |
| p115D | PX069869 | PX069887 | PX069902 |
| p1236 | PX101737 | PX095409 | PZ568904 |
| p1239 | PX101738 | PX095410 | PX111620 |
| p1241 | PX101739 | PX095411 | PX111621 |
| p1242 | PX098497 | PX098500 | PX098503 |
| p126 | PX069871 | PX069889 | PX069903 |
| p127 | PX069872 | PX069890 | PX069904 |
| p1313 | PX098498 | PX098501 | PX098504 |
| p1316 | PX098921 | PX098927 | PX098933 |
| p133 | PX069874 | PX069892 | PX069905 |
| p136 | PV796316 | PV796317 | PV796318 |

| | | | |
|---|---|---|---|
| p137 | PV808467 | PV808468 | PV808469 |
| p1427 | PX098924 | PX098930 | PX098934 |
| p1430 | PX098925 | PX098931 | PX098935 |
| p1918 | PX101724 | PX095415 | PX111625 |
| p1921 | PX101725 | PX095416 | PX111619 |
| p201 | PX101730 | PX095392 | PX111627 |
| p332 | PX069876 | PX069894 | PX069906 |
| p334 | PX069877 | PX069895 | PX069907 |
| p502 | PX069880 | PX069898 | PX069908 |
| p915 | PX069881 | PX069899 | PX069909 |
| p931 | PX069883 | PX069901 | PX069910 |
| Spain-Case 10 | PP 00702S6.3 | PP 00702S6.3 | PP 00702S6.3 |
| Spain-Case 12 | PP 0072Q1X.1 | PP 0072Q1X.1 | PP 0072Q1X.1 |
| Switzerland-Case 7 | PP 006WBLH.2 | PP 006WBLH.2 | PP 006WBLH.2 |

**Supplementary Methods**

***Detailed Bayesian phylogenetic analyses***

Bayesian phylogenetic analyses were conducted independently for the S, M, and L genomic segment datasets using BEAST v1.10.5. Analyses implemented a codon-partitioned scheme separating codon positions 1+2 from position 3. Independent GTR+Γ substitution models were assigned to each partition, with empirical base frequencies estimated independently and among-site rate heterogeneity modeled using a discrete gamma distribution with four categories.

A strict molecular clock model was applied without temporal calibration or incorporation of sampling dates, such that branch lengths represented relative genetic divergence rather than evolutionary time. Tree inference was performed under a constant-size coalescent prior.

Initial unconstrained analyses were conducted to evaluate the natural phylogenetic placement of outbreak-associated genomes within broader South American Andes virus diversity. Additional analyses incorporated a monophyletic constraint for outbreak-associated genomes to evaluate clustering stability across genomic segments.

Markov chain Monte Carlo (MCMC) chains were run for 200 million generations with automatic operator optimization enabled. Tree-space exploration used standard BEAST topology operators including subtree-slide, narrow exchange, wide exchange, and Wilson–Balding moves together with scale and delta-exchange operators applied to substitution model, frequency, gamma-shape, and coalescent parameters.

Maximum clade credibility (MCC) trees were generated following removal of burn-in and visualized using FigTree v1.4.4. Posterior support values associated with major phylogenetic nodes were incorporated into the final tree visualizations.

***Detailed phylogeographic analyses***

Discrete phylogeographic reconstruction was carried out independently for the S, M, and L segments in BEAST 10.5.0, using identical configurations across segments. Each dataset (sampling years 1996–2019) was analyzed under a single GTR+Γ substitution model (four categories) with substitution parameters, base frequencies, and the gamma shape estimated. Tips were calibrated by year of sampling under a strict molecular clock, with the clock rate estimated under a CTMC-scale reference prior and a constant-size coalescent tree prior. Temporal signal in these datasets was weak, as assessed by root-to-tip regression in TempEst v1.5.3 ($R^2 = 0.32$); absolute divergence times and the timing of individual geographic transitions were therefore treated as unreliable and not interpreted. Because discrete ancestral-state reconstruction and the identification of supported transitions depend on tree topology and the relative distribution of character changes rather than on absolute calibration, phylogeographic inference was restricted to the identity and posterior support of geographic transitions. Each genome sequence was assigned a discrete geographic location corresponding to the most probable site of infection from epidemiological investigation, comprising 15 states across administrative regions of southern Chile and localities of northern Patagonia (Argentina) and including an explicit "unknown" category for genome sequences without a confidently inferred infection site. Ancestral locations were reconstructed under a symmetric continuous-time Markov chain model (all pairwise exchange rates estimated), and transitions among locations were quantified by Markov jump counting with complete-history logging. Well-supported transitions were defined as those present in >80% of the posterior trees (posterior probability > 0.80).

Convergence and mixing were assessed in Tracer v1.7.2; after discarding the first 10% of each chain as burn-in, all estimated parameters had effective sample sizes (ESS) above 200. Maximum clade credibility trees were generated in TreeAnnotator v10.5.0 and visualized in iTOL v4. Well-supported geographic transitions were mapped in QGIS v3.44 for visualization.

***Detailed reassortment analysis***

To test whether pairwise genetic distances were concordant across genomic segments, all pairwise p-distances were computed among Clade III viruses for the S, M, and L segments. For each isolate pair and each segment comparison $(a,\ b) \in \{(S, M), (S, L), (M, L)\}$, the number of observed nucleotide differences $(Y_a,\ Y_b)$ and the number of comparable sites $(L_a,\ L_b)$ were recorded.

Substitution counts were modelled as independent Poisson random variables sharing an isolate-pair rate $r$ and a constant segment rate ratio $\lambda$:

$$Y_a \sim Poisson(L_a r), \qquad Y_b \sim Poisson(L_b \lambda r).$$

Conditioning on the total number of differences $T = Y_a + Y_b$ eliminates $r$ and yields

$$Y_b \mid T \sim Binomial(T, \pi), \qquad \pi = \lambda L_b \,/\, (L_a + \lambda L_b).$$

The rate ratio $\lambda$ was estimated by conditional binomial maximum likelihood from all pairs with expected $Y_b$ count $T_\pi > 20$, and the fitted relationship was displayed on the p-distance scatter as the line $y = \lambda x$. For each pair meeting the same $T_\pi > 20$ filter, departure from the null expectation under the fitted $\lambda$ was assessed with a two-sided normal approximation,

$$Z = \frac{Y_b - T_\pi}{\sqrt{T\pi(1-\pi)}} \sim N(0,1),$$

Within each segment comparison, p values were corrected for multiple testing by the Benjamini–Hochberg procedure, and pairs with q < 0.05 were considered significant.

To visualize local variation in genetic distance along each segment, pairwise p-distances were calculated in a sliding window of 100 consecutive comparable sites with a stride of 10 comparable sites for selected isolate pairs. KS statistics was applied to test whether the nucleotide differences appear uniformly across the genome.

***Detailed structural modeling of the Andes virus L protein core domain***

The core domain of the ANDV L protein was generated using the SWISS-MODEL web server[8]. The coding sequence of the Chile-9717869 isolate was used as the query, and the crystal structure of the Sin Nombre virus L protein bound to the 5' viral RNA (PDB: 8CI5)[5] was selected as the template. The target and template shared 86,3% sequence identity. The resulting model comprised a monomer spanning residues 225 to 1602 of the L protein. Model quality was assessed using the QMEANDisCo global score, leveraging a score of 0.71 ± 0.05 (scale 0 to 1), indicating moderate-to-high confidence in the predicted structure. Structural superposition analyses were performed using both the SNV L protein structure (PDB: 8CI5)[5] and the hexameric Hantaan virus polymerase structure (PDB: 8QHD)[9] as references, leading to a root-mean-square deviation (RMSD) of 0.189 Å for SNV L, while a RMSD of 1.869 Å was obtained for HTNV L, consistent with greater structural divergence.

***Structural coordinate superposition and local interaction analyses***

Detailed structural analyses were performed using experimentally determined and Alpha Fold 3-predicted ANDV protein models. Structural coordinate visualization, residue mapping, and local interaction analyses were conducted in UCSF ChimeraX v1.11. Structural analyses included the native $G_N/G_C$ spike tetramer (PDB ID: 9P3X)[1] the $G_C$ post-fusion trimer (PDB ID: 6Y6Q)[3], the $G_N$ zinc-finger domain (PDB ID: 2K9H)[2], and the L protein N-terminus variant K124A (PDB ID: 6Q99)[4]. Additional structural contextualization was performed using Alpha Fold 3-predicted models for unresolved $G_N/G_C$ regions[10] and the ANDV L protein core domain. Alpha Fold 3-predicted models were structurally superimposed onto experimentally determined coordinates using the MatchMaker extension implemented in ChimeraX under the BLOSUM-62 matrix and a 2.0-Å pruning cutoff. Structural similarity between models was evaluated using root-mean-square deviation (RMSD) analyses and aligned residue comparisons.

Predicted effects of naturally occurring amino acid substitutions on protein stability and intermolecular interactions were evaluated using DDMut and DDMut-PPI[11,12]. DDMut analyses were applied to monomeric domains, whereas DDMut-PPI was used for oligomeric assemblies including $G_N/G_C$ tetramers and protomer interaction interfaces. For *N*-glycosylations on $G_N/G_C$, the effect of SNVs was screened for the amino acid motive NXS/T.

#### Supplementary Results

##### *Supplementary phylogenetic analyses support placement of outbreak-associated genomes within the HHPC cluster embedded in Clade III Andes virus diversity*

Maximum clade credibility trees inferred from Bayesian phylogenetic analyses of the S, M, and L genomic segment sequences yielded topologies consistent with the maximum-likelihood phylogenies (**Supplementary Fig. S1**). Across all three genomic segments, outbreak-associated genome sequences were assigned to the Chilean sequence-enriched cluster, the HHPC cluster, embedded in ANDV Clade III previously described for southern South American ANDV diversity and grouped with historical Chilean and northern Patagonian genome sequences.

The overall topological structure of the S, M, and L segment trees was concordant across phylogenetic inference approaches. Similar clustering patterns were observed independently for sequences of each genomic segment, including grouping with genome sequences from Los Ríos and Araucanía Regions in southern Chile together with related Patagonian genomes from Argentina.

Because outbreak-associated genome sequences were minimally divergent, a single representative genome sequence was used in the main phylogenetic analyses.

##### *Genome-wide analyses provide little evidence of reassortment within Andes virus Clade III*

To test whether there is reassortment in these viruses, we built the p-distance test based on the hypothesis that without segment exchange, the nucleotide substitutions accumulate in the genome randomly. As a result, pairwise distances on S, M, and L should remain proportional (a roughly constant segment rate ratio). Overall, pairwise distances among Clade III segments are largely concordant, indicating few reassortment events have occur in these viruses (**Supplementary Figure S2a–c**). The outliers involve p1059 and p1236. For p1059, S is nearly identical to NRC-4/18 while M and L are not, which could indicate recent shared S ancestry (**Supplementary Figure S2d**), although the small number of S differences warrants cautious interpretation given possible artifacts. For p1236, elevated L distance is concentrated in the second half of the segment (KS $D = 0.182$, $p = 3.29 \times 10^{-3}$; **Supplementary Figure S2e**). Although this localized pattern could be consistent with recombination, homologous recombination is considered uncommon in negative-sense RNA viruses and was not supported by the broader phylogenetic evidence. Together, these results provide little evidence that reassortment has shaped Clade III genome relationships.

***Detailed structural characterization of enriched substitutions identified within the Hua Hum Pass corridor cluster***

The three HHPC-enriched SNVs resulting in changes within $G_N/G_C$ ectodomains (I114V, A193T, and S1055T) map to surface-exposed regions of the tetrameric spike structure (**Supplementary Figure S3a–e**). I114V is positioned at the $G_N$ head domain, whereas A193T lies on its lateral surface, close to the central groove formed by the $G_N$ tetramer (**Supplementary Figure S3b–c**). S1055T is exclusively encoded by two HHPC sequences from Villa Meliquina, Province of Neuquén, Argentina. The change introduces a novel *N*-glycosylation motive that is not encoded by cruise ship outbreak viruses, nor other HHPC viruses or ANDV linages in general (**Supplementary Table S3**). Given its proximity to the center of the tetrameric spike, this glycosylation site could influence the accessibility of this region. In contrast, S1055T is exposed on the surface of $G_C$ domain III (DIII), where it is accessible only when the lateral face of $G_N/G_C$ is not engaged in lattice contacts with neighboring tetramers (**Supplementary Figure S3d**). The same residue is also exposed on the surface of $G_C$ DIII in the post-fusion homotrimer (**Supplementary Figure S3i**).

The remaining two HHPC-associated GPC substitutions are within region that is not accessible to antibodies. As previously reported[10], the outbreak-specific change $G_N$ T516I maps to a region that remains unresolved in available experimental structures, and is expected to lie at the membrane–water interface, forming either part of $G_N$ transmembrane domain 1 or of the $G_N$ endodomain (**Supplementary Figure S3g,h**). The HHPC-specific $G_C$ V1127I substitution corresponds to the last residue resolved within the $G_C$ transmembrane domain in the native tetramer structure[13] (**Supplementary Figure S3f**).

Within L protein, R144K maps to the endonuclease domain, but is positioned away from the catalytic center involving $Mn^{2+}$[4] (**Supplementary Figure S4a**). A541V was assessed using the Alpha Fold-3 model generated from the L protein core structure (i.e., the region containing the core lobe, vRNA-binding lobe, and the canonical fingers, palm, thumb, bridge, and thumb-ring domains)[5]. The position is solvent-exposed and is part of the viral

RNA-binding lobe where it contacts I544 (**Supplementary Figure S4b–c**). Both substitutions, R144K and A541V, are conservative and are not predicted to perturb the local structural environment.

The effects of individual and combined substitutions on protein stability and inter-heterodimeric affinity within the $G_N/G_C$ tetramer and on the L protein endonuclease and core domains were evaluated using deep learning-based predictors DDMut and DDMutPPI[12].

*In silico* analyses revealed only minor changes in Gibbs free energy (ΔΔG), indicating that these substitutions are unlikely to substantially affect the structural homeostasis of the proteins. The limited structural impact is consistent with the conservative physiochemical nature of the residue substitutions, which preserve the key properties of native residues and therefore are not expected to perturb the tertiary or quaternary organization of the proteins
